# Integrating Single-Cell Biophysical and Transcriptomic Features to Resolve Functional Heterogeneity in Mantle Cell Lymphoma

**DOI:** 10.1101/2025.05.20.655210

**Authors:** Ye Zhang, Lydie Debaize, Adam Langenbucher, Jenalyn Weekes, Ioulia Vogiatzi, Teemu P. Miettinen, Mingzeng Zhang, Emily Sumpena, Huiyun Liu, Sarah M. Duquette, Liam Hackett, Jeremy Zhang, Sona Baghiyan, Robert A. Redd, Martin Aryee, Matthew S. Davids, Austin I. Kim, Christine E. Ryan, David M. Weinstock, Scott R. Manalis, Mark A. Murakami

## Abstract

Intra-tumor heterogeneity impacts disease progression and therapeutic resistance but remains poorly characterized by conventional histologic, immunophenotypic, and molecular approaches. Single-cell biophysical properties distinguish functional phenotypes complementary to these approaches, providing additional insight into cellular diversity. Here we link both buoyant mass and stiffness to gene expression to identify clinically relevant phenotypes within primary mantle cell lymphoma (MCL) cells, employing MCL as a model of biological and clinical diversity in human cancer. Linked measurements reveal that buoyant mass and stiffness characterize B-cell development states from naïve to plasma cell and correlate with expression of oncogenic B-cell receptor signaling genes such as *BLK* and *CD79A*. Additionally, changes in cell buoyant mass within primary patient specimens *ex vivo* correlate with sensitivity to Bruton’s Tyrosine Kinase inhibitors *in vivo* in MCL and chronic lymphocytic leukemia, another B-cell malignancy. These findings highlight the value of biophysical properties as biomarkers of response in pursuit of future precision therapeutic strategies.

## INTRODUCTION

Biophysical parameters, such as cell buoyant mass and stiffness, are quantitative measures that integrate a wide array of molecular components and cellular processes, including metabolism, growth, and cytoskeletal dynamics (*1–6*). In contrast, molecular data, such as gene expression profiles, offer detailed insights into specific pathways and processes, providing mechanistic insights into cellular phenotypes. Thus, the two approaches are potentially complementary. Biophysical parameters have a practical advantage in some settings, as they can be easier and faster to measure than high-content molecular assays (*7*, *8*). If a clinically meaningful molecular state can be reliably inferred from a simpler biophysical measurement, it could offer a more accessible and efficient platform for translating these insights into diagnostic or therapeutic biomarker strategies.

Previous studies have provided initial proof-of-concept for linking biophysical properties with molecular data. For example, Xu et al. measured the average stiffness of cell lines and correlated these measurements with gene expression profiles, revealing associations between mechanical properties and specific molecular pathways related to metastatic potential (*9*). While this approach offers a broader understanding of how cellular stiffness is influenced by underlying genetic programs, averaging measurements across entire populations can obscure cell-to-cell variability. Addressing this, Kimmerling et al. linked individual cell buoyant mass directly to gene expression profiles at the single-cell level. This approach revealed subtle relationships related to cell cycle progression that were inapparent in population-level studies (*10*). Despite these advances, a major limitation of these studies is that they have been performed primarily in cell lines rather than in primary cells, which do not fully represent the complexity and variability of cells found in primary tissues, including crucial microenvironmental interactions.

To enhance the translational relevance of linked biophysical and transcriptional measurements, we examined their functional correlations in models that more accurately reflect the biological complexity of in situ disease. Central to this fidelity is inter- and intra-tumor heterogeneity in cell type composition and tumor cell genomics, transcriptomics, and oncogenic signaling. To capture this complexity, we employed a diverse spectrum of complementary cancer model systems, including cell lines, patient-derived xenografts (PDXs), and primary patient tumor specimens, to: i) define the range of lineage-specific developmental biophysical phenotypes, ii) link biophysical and transcriptional phenotypes of individual cells, and iii) establish their associations with patient clinical outcomes. As a clinically relevant setting for a novel high-resolution diagnostic platform, we investigated the translational potential of linked multiparametric biophysical and transcriptional measurements in mantle cell lymphoma (MCL) at the single-cell level.

MCL, which is a subtype of B cell non-Hodgkin lymphoma that exhibits both aggressive and indolent clinical features (*11*), is an archetype of cancer biological and clinical heterogeneity (*12*). It encompasses divergent cell states that impact disease progression and response to treatment. On one end of this clinical spectrum are patients with highly indolent disease that can be observed after diagnosis. In particular, the leukemic non-nodal variant of MCL is often diagnosed as an incidental finding in patients who have no symptoms (*13–16*). On the other end of the spectrum are patients who experience a much more aggressive course that mimics high-grade lymphomas (*17*). There are some well-established correlates of risk, including histologic features (i.e. blastoid, pleomorphic), clinical characteristics (as included in the MCL International Prognostic Index [MIPI] score)(*18*), and genomic features (i.e. *TP53*- aberrancy) (*16*, *19–21*). However, histopathologically and molecularly diverse subclones may co-exist. In addition, risk stratification methods using these factors are conventionally applied at diagnosis, but lymphomas can evolve over time, typically from a more indolent to a more aggressive disease state. A facile approach to better resolve MCL heterogeneity at each stage of the disease course could guide precision therapeutic strategies optimally aligned with individual patient risk and disease biology. By analyzing primary MCL cells, we aimed to uncover how biophysical properties like cell buoyant mass and stiffness correlate with molecular features and illuminate functional disease heterogeneity.

## RESULTS

We previously developed the Suspended Microchannel Resonator (SMR)(*22*) to measure the buoyant mass of single cells. The SMR achieves more than ten times the precision of traditional microscopy for measuring cell size, providing new insights into cellular growth and response to perturbations. In addition to its high measurement precision, buoyant mass is a unique biophysical parameter because it represents the product of a cell’s volume and its buoyant density relative to the surrounding fluid. For example, we recently demonstrated that in primary T cells, buoyant mass reveals cell states that are not detectable from volume or buoyant density alone (*23*). Recognizing that both buoyant mass (hereafter referred to simply as mass) and stiffness may uncover unique aspects of cell state (Fig. S1A-C)(*24*), we leveraged the SMR device to measure both properties within the same cell. Analysis of six MCL cell lines, Jeko-1, Granta-519, Maver-1, Mino, Rec-1 and Z-138, revealed distinct single-cell mass and stiffness profiles (Fig. S1D, E). While these cell lines display measurable variability, patient-derived xenografts (PDXs) capture more of the cellular diversity and complexity of MCL patients, providing greater relevance for studying therapeutic responses. To investigate the correlation between these biophysical properties and the morphologic, immunophenotypic, and clinical characteristics of MCL, we examined three PDX models of MCL from the Public Repository of Xenografts (*25*) that encompass a spectrum of disease genetics and growth kinetics. These included PDX DFBL-96069, derived from a patient with classic MCL, and DFBL-39435 and DFBL-91438, from patients with blastoid and pleomorphic variants, respectively, each with unique mutational profiles, more aggressive clinical behavior and adverse prognoses (Table 1 and Table S1) (*19*, *26*).

**Table 1.** Clinical and demographic summary of MCL PDX patients

| PDX model | Patient sex-age (years) | Organ of origin | Patient clinical summary | Patient cytogenetics | PDX histologic subtype |
| --- | --- | --- | --- | --- | --- |
| DFBL-96069 | F-72 | PB | Relapsed/Refractory MCL following several lines of chemotherapy (R-CHOP-like regimen* +Et/Cyt, COPD-L, FC, Teniposide+ Cytarabine, Cladribine, Lenalidomide+ Dexamethasone). Refractory and intolerant to recent ibrutinib. | del(6)(q13q23), del(13)(q12q22) t(11;14)(q13;q32) indicating the presence of the *CCND1/IGH* translocation. | Classic |
| DFBL-39435 | M-67 | PB | Relapsed/Refractory MCL after Rituximab-Bendamustine and Rituximab maintenance; then Bortezomib | 11q13(*CCND1*x3),14q32(*IGH*x3),(*CCND1* con *IGH*x2),17p13.1(*TP53* x1) indicating the presence of the *CCND1/IGH* translocation and a *TP53* deletion. | Blastoid |
| DFBL-91438 | M-75 | PB | Relapsed/Refractory MCL following several lines of chemotherapy (R-CHOP, R-CVP, FMR Zevalin, Bortezomib + Decadron + Rituxan). Complete remission to Palbociclib | Not done | Pleomorphic |

C: cyclophosphamide, Cyt: cytarabine, D: daunorubicin, Et: etoposide/teniposide, F: fludarabine, H: doxorubicin hydrochloride (hydroxydaunomycin), L: L-asparaginase, M: mitoxantrone, O or V: oncovin/vincristine sulfate, P: prednisone, R: rituximab, \*: regimen includes cyclophosphamide, vindesine, epirubicin (used as anthracycline), and prednisone.

PDX cells were harvested from spleens of engrafted mice and underwent comprehensive biophysical analysis using the SMR, along with immunophenotyping by flow cytometry, histopathological examination, and bulk RNA sequencing (Fig. 1A). Concordant with clinical hematopathology evaluation of the patient specimens from which they were derived, all PDX tumor cells were positive for CD45, CD19, CD5, and monotypic surface immunoglobulin light chain by flow cytometry (Fig. S2A). Immunohistochemistry (IHC) showed strong expression of cyclin D1 and surface CD20 in tumor cells from all three models, while DFBL-39435 also showed overexpression of p53 (Fig.1B and Fig. S2B). Whole exome sequencing (WES) revealed mutations in genes recurrently altered in MCL including *KMT2C*, *ATM*, and *RB1* for DFBL-96069, *TP53*, *KMT2C*, *DNMT3A* and *MYC* for DFBL-39435, and *TP53* for DFBL-91438 (Table S1 for the full list). All three models exhibited diffuse growth patterns and pleomorphic features with irregular nuclei, prominent nucleoli, high mitotic activity, and elevated Ki67 proliferation indices (Fig.1B). Enrichment of these histomorphological characteristics, which have been associated with aggressive clinical behavior (*26*, *27*), is a common phenomenon following xenotransplantation of MCL.

**Figure 1.**
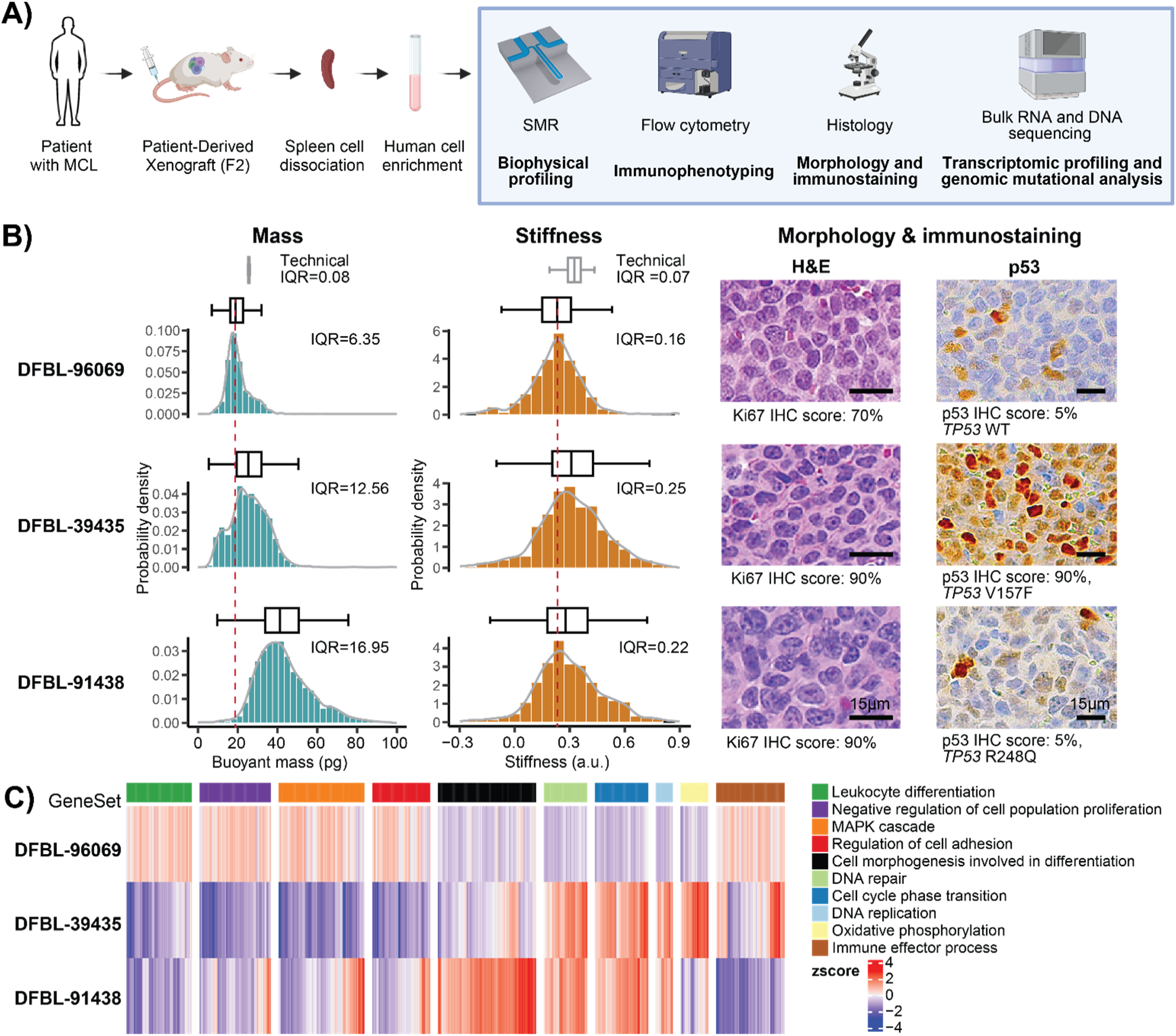
Comparative analysis of biophysical properties, morphological attributes, and transcriptomic features in primary human MCL cells. **(A)** Schematic workflow for isolating tumor cells from MCL PDX models for biophysical profiling, immunophenotyping, histopathologic imaging and bulk RNA/DNA sequencing. Created with BioRender. **(B)** Histogram distributions of single-cell buoyant mass and stiffness measurements using the SMR, along with representative H&E stain and p53 immunohistochemistry (IHC) images (40X, scale bar: 15 µm) of human-enriched cells from the spleen of three PDX models: DFBL-96069, DFBL-39435, DFBL-91438. Mass and stiffness profiles are from one representative replicate of 3 repeats. A box plot representing the median and interquartile range (IQR), is shown above each histogram, with the median mass and stiffness of DFBL-96069 indicated by a dotted red line for reference. The technical IQR of mass was determined by repeatedly measuring the same 10 μm polystyrene bead using the SMR and that of stiffness by repeatedly measuring the same mouse lymphocytic leukemia cell line L1210 (Fig. S3A-E). The *TP53* mutational status based on targeted sequencing (Table S1) and the relative percentage of p53 and Ki67 positive cells based on IHC stains (Fig. S2B) are described below the H&E and IHC images. **(C)** Gene set signature enrichment evaluation of bulk RNA sequencing of MCL cells isolated from the spleen of the three PDXs.

Biophysical profiling unveiled marked differences in mass and cellular stiffness between and within tumor samples, highlighting inter- and intra-tumor heterogeneity. MCL cells from DFBL-39435 and DFBL-91438 exhibited a higher mass and increased stiffness compared to those from DFBL-96069 (Fig. 1B). Notably, the observed variability was biological, as it greatly exceeds the technical noise (technical interquartile ranges, IQRs) of our measurements (Fig. S3)(*28*). Given that mass reflects cell density and volume, we utilized a fluorescence exclusion-coupled suspended microchannel resonator (fxSMR)(*29*), an established method for measuring these additional biophysical characteristics. This analysis showed that MCL cells from DFBL-91438 model (median size 1080 µm^3^) and DFBL-39435 (median size 859 µm^3^) were larger than those from DFBL-96069 (median size 794 µm^3^). We also found that MCL cells from DFBL-91438 and DFBL-39435 had higher cell density compared with those from DFBL-96069 (Fig. S2C). Using Amnis® Imaging Flow Cytometry, we also evaluated cell surface area and perimeter length. We found that DFBL-91438 cells had the largest area and longest perimeter, followed by DFBL-39435 and DFBL-96069 (Fig. S2C). These findings orthogonally corroborated the phenotypic variability across MCL subtypes that we detected with the SMR. Additionally, bulk RNA-sequencing of the three PDX models revealed differences across models in the expression of genes involved in pivotal pathways, including leukocyte differentiation, cell adhesion, MAP kinase signaling, oxidative phosphorylation (OXPHOS), and cellular differentiation (Fig. 1C). Notably, the DFBL-96069 model exhibited lower expression of genes critical for cell cycle progression and DNA replication, along with higher expression of genes that act as proliferation inhibitors, compared to DFBL-39435 and DFBL-91438, consistent with Ki67 levels observed by IHC (Fig. 1B). These findings not only underscore the biophysical variability across MCL subtypes but also highlight transcriptional differences, raising important questions about how these transcriptional profiles correlate with biophysical properties at the single-cell level.

### Paired single-cell biophysical and transcriptional characterization of MCL

Cell mass and stiffness are key biophysical properties driven by distinct molecular and transcriptional cell states. To investigate these relationships, we developed a single-cell SMR-RNA sequencing (scSMR- RNA-Seq) platform to integrate individual biophysical and transcriptional measurements at single-cell resolution (Fig. 2A). Cells were isolated after biophysical measurement on the SMR device and collected into a PCR-tube strip on a motorized stage, at a throughput of up to 120 cells per hour. Each cell then underwent scRNA-Seq employing the Smart-seq2 protocol, which generates deep transcriptome profiles of individual cells (average depth of 0.5-1 million reads per cell, detecting up to 6,000-10,000 genes per cell). Smart-seq2 offers high sensitivity for low-abundance transcripts and full-length transcript coverage, enabling comprehensive gene expression profiling (*30*, *31*). Using this workflow, human MCL cells were harvested from engrafted PDX models in NOD.SCID.IL2Rγ^-/-^ (NSG) mice, purified from various tissues, including bone marrow, liver, peripheral blood, and spleen, and subjected to linked biophysical- transcriptomic profiling (Fig. S4A-E). This integrative approach provides a framework for linking biophysical properties with functionally distinct states at single-cell resolution.

**Figure 2.**
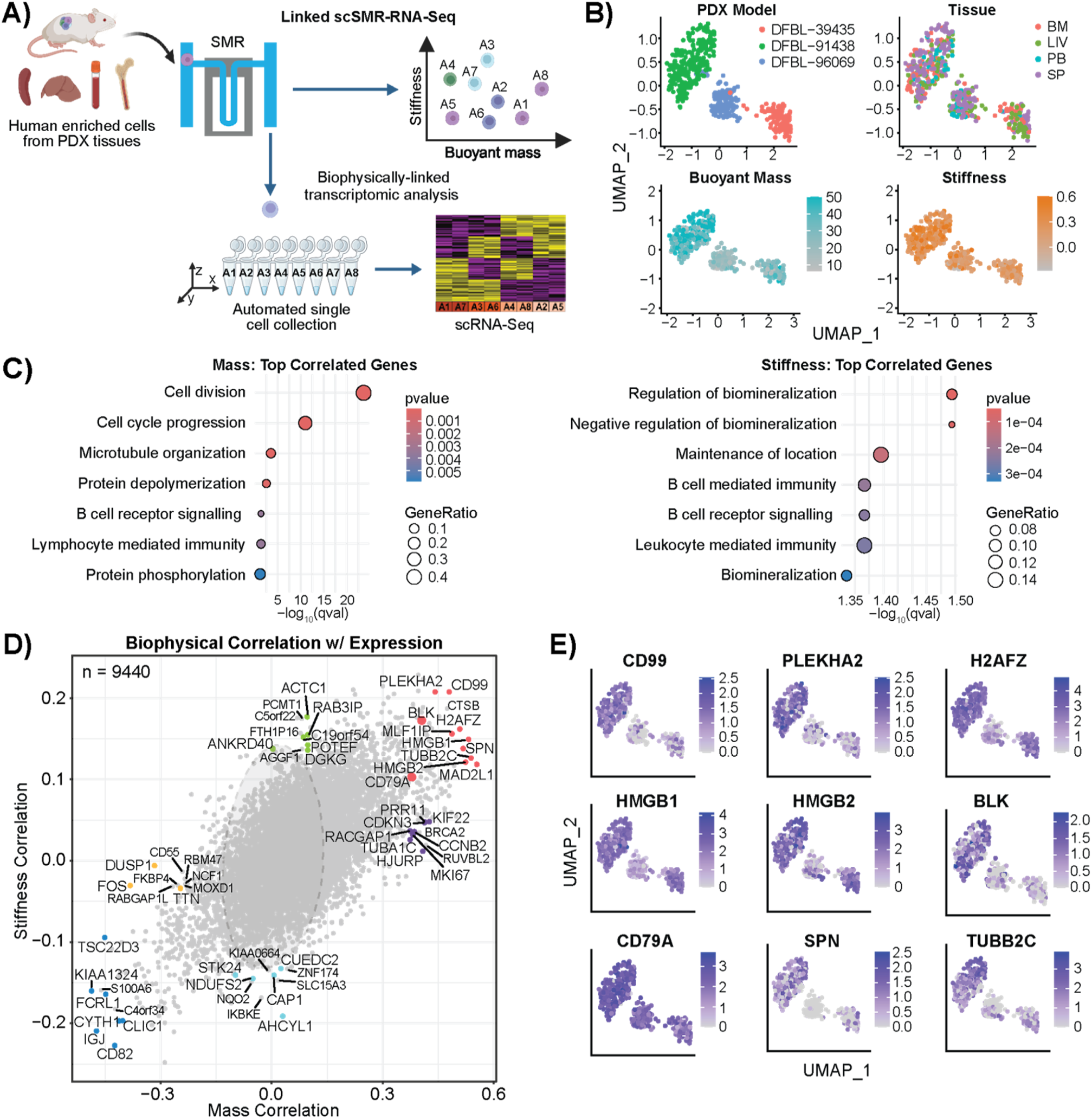
Cell mass and stiffness are strongly associated with the expression of genes annotated for cell division, cell cycle, and B-cell activation. **(A)** Schematic workflow linking the biophysical measurements of individual MCL cells from different tissues of three PDX models to downstream scRNA-Seq. Created with BioRender. **(B)** UMAP analysis of linked single-cell mass, stiffness and scRNA-Seq data of human tumor cells enriched from DFBL-96069, DFBL-39435, DFBL-91438 isolated from different tissues (SP, spleen; PB, peripheral blood; BM, bone marrow; and LIV, liver). Biological replicates, each from a different mouse, were included per tissue, as detailed in Fig. S4. **(C)** GSEA of the top genes correlated with mass (left) and stiffness (right) measurements among the combined PDX cells. The color represents the *p*-values, and the size of the spots represents the gene ratio, which is defined as the number of genes in the overlap divided by the total size of the gene set. **(D)** Biophysical correlation with gene expression in MCL PDXs. Representative genes are color-coded as follows: red for genes positively correlated with both mass and stiffness, purple for genes positively correlated with mass only, green for genes positively correlated with stiffness only, dark blue for genes negatively correlated with mass and stiffness, light blue for genes negatively correlated with stiffness only, and yellow for genes negatively correlated with mass only. **(E)** UMAP plots showing the expression of nine representative genes positively correlated with MCL cell mass and stiffness, marked as red in Fig. 2D.

Uniform Manifold Approximation and Projection (UMAP) visualization of transcriptomes revealed that cell clustering was determined more by PDX model than tissue of origin (Fisher’s Exact test, *p*-value < 2.2e- 16, Fig. 2B). Notably, single-cell mass and stiffness were not correlated across the three PDX models or within each individual model, as indicated by low R² values (0.0025, 0.002, 0.001, and 0.0013 respectively; Fig. S5A). This finding highlights the necessity of measuring both mass and stiffness as orthogonal features. It also motivates further investigation into how distinct transcriptional factors regulate these biophysical properties. Transcriptome-wide correlation analyses within each PDX model identified three shared genes, *CKS1B, TUBB,* and *DEPDC1B*, among the top 100 most correlated with cell mass (Fig. S5B). These shared genes are known to play a key role in cell cycle regulation (*32–34*), suggesting a tight association between cell cycle progression and cell mass. Interestingly, DFBL-96069 displayed a higher proportion of cells in the G0/G1 phase, while DFBL-39435 and DFBL-91438 were enriched for cells in the G2/M phase (Fig. S6A), consistent with earlier bulk-sequencing results (Fig. 1C). Cells in the G2/M phase from all three models exhibited higher mass compared to those in the G0/G1 and S phases, reflecting the increased cellular content associated with mitosis (Fig. S6B). Additionally, two MCL cell lines, Rec1 and Jeko-1, displayed approximately 30% and 45% greater mass compared to controls, respectively (Fig. S6C-D), when synchronized in G2/M phase using nocodazole treatment, functionally confirming the expected relationship between cell cycle progression and cell mass in MCL. These findings extend similar observations previously reported in acute leukemias (*4*, *10*, *35*). However, cell cycle distribution only partially explains the mass variation observed across PDX models. For instance, cells from DFBL-91438 in the G2/M phase had a higher mass compared to those in G2/M from DFBL- 96069 and DFBL-39435. Similar results were observed from cells in S and G0/G1 phases (Fig. S6B). These data suggest that additional molecular programs beyond those controlling cell cycle progression contribute to inter-model cell mass differences.

To determine the molecular programs most correlated with biophysical properties across models, we performed Gene Set Enrichment Analysis (GSEA) among the three PDX models. To focus on biologically relevant differences rather than inherent model-specific factors, such as genes linked to sex or immunoglobulin expression, we excluded model-specific genes (i.e. those expressed in less than 10% of cells in any model). Pathways annotated for cell division, cell cycle progression, microtubule organization, chromosome segregation, and cell proliferation (Fig. 2C and Table S2) were enriched with top genes (z- score>2.5, Fig. S4F) correlated with mass. In addition to these cell division and growth-related pathways, which comprise the majority of the top 100 hits, we identified other significant enrichment (*q*-value<0.1, Fig. 2C) of pathways involved in B-cell receptor (BCR) signaling and protein phosphorylation. The top 10 genes correlated specifically with mass (Fig. 2D, marked in purple) include *CDKN3, CCNB2* and *RACGAP1* (critical for cell cycle progression), *MKI67* and *PRR11* (proliferation markers), and *KIF22, HJURP* and *TUBA1C* (involved in mitotic processes). In contrast, *DUSP1* and *FOS*, which are negatively correlated with cell mass (Fig. 2D, marked in orange), play key roles in cellular stress responses and survival (*36*, *37*). The top genes (z-score>2.5, Fig. S4F) correlated with cell stiffness were enriched in pathways linked to the adaptive immune response and BCR signaling (Fig. 2C), including *BLK*, *CD99, IRF4, SPIB, FOXP1*, *SWAP70* and *TCL1A* (Table S3). Furthermore, the top 10 genes specifically correlated with cell stiffness but not mass (Fig. 2D, marked in green) included *ACTC1, POTEF* and *ANKRD40* (linked to cytoskeletal organization), *C19orf54* (actin maturation), *AGGF1* (adhesion and extracellular interactions), and *RAB3IP* and *DGKG* (involved in trafficking and lipid signaling).

Additionally, transcriptome-wide correlation analysis further identified subsets of genes positively associated with both mass and stiffness (Fig. 2D, marked in red). Among the top correlated genes were *CCND1*, a key regulator of cyclin-dependent kinase activity and cell cycle progression translocated in all three models to the *IGH* locus (*38*, *39*); *HMGB1/HMGB2*, chromatin structural factors involved in transcriptional regulation, DNA replication, and repair (*40*, *41*); *MLF1IP*, which is required for centromere assembly and promotes cancer cell proliferation by regulating mitotic progression and genome stability (*42*); the histone variant *H2AFZ*, which is critical for chromatin structure, chromosome segregation, cell cycle progression and has been linked to tumor progression in various cancers (*43*, *44*); as well as *PLEKHA2*, which is implicated in the positive regulation of cell-matrix adhesion (*45*). Together, these genes provide mechanistic insight into the biophysical traits that integrate proliferation, actin dynamics, and cell adhesion (*3*, *9*, *10*). Genes involved in BCR signaling, including *BLK* (B-lymphoid tyrosine kinase) and *CD79A*, positively correlated with both mass and stiffness. Genes involved in immune response and leukocyte migration either positively (*SPN, HLA-DPA1*, *HLA-DMA HLA-DRA*, and *CD99*) or negatively (*FCRL1, IGJ*, *CYTH1*, and *CD82*) correlated with both mass and stiffness (Fig. 2D, marked in dark blue and Table S3). Additionally, genes positively associated with both mass and stiffness that contribute to DNA repair or DNA damage response included *BCLAF1*, *H2AFX*, *MAD2L1*, *POLD1*, and *TP63*. These findings highlight the capability of paired biophysical-transcriptomic characterization to link biophysical properties to broadly generalizable as well as MCL-specific molecular programs. Moreover, they support the hypothesis that cell mass and stiffness reflect distinct molecular programs governing not just cell growth and proliferation but also pathways with oncogenic relevance.

Genes that are positively correlated with both cell mass and stiffness in Fig. 2D (marked in red) appear to be primarily driven by inter-model differences rather than a direct correlation between these biophysical properties at the single-cell level. Specifically, cells from DFBL-91438 exhibited the highest mass and stiffness, whereas those from DFBL-96069 had lower stiffness and lower mass. The spatial distribution of top correlated genes within the UMAP plot further supported these findings (Fig. 2E). For example, genes such as *CD99* and *SPN*, as well as the BCR signaling gene *BLK*, were predominantly expressed in the proliferative models DFBL-39435 and DFBL-91438 (Fig. 2E). These findings show how model-specific differences in cellular and molecular states can drive observed biophysical variation, underscoring the need to account for both inter- and intra-tumor heterogeneity when interpreting the relationship between biophysical properties and transcriptional profiles.

### Biophysical characterization of naïve B cell activation

Our paired analysis, which identified the well-established correlation between cell mass and cell cycle (*4*, *10*, *35*), also revealed an association of both mass and stiffness with the BCR signaling pathway. This pathway plays a central role in the pathogenesis of MCL by driving cell proliferation, survival, and therapeutic resistance (*46*). Therefore, we decided to further examine the influence of B-cell activation on biophysical properties. We profiled single-cell biophysical characteristics across the spectrum of normal antigen-dependent B-cell development, from naïve B-cells to plasma cells. Naïve B cells were isolated from peripheral blood mononuclear cells (PBMCs) of three healthy donors via immunomagnetic separation. These cells were stimulated *ex vivo* to trigger activation and promote differentiation (Fig. 3A). *Ex vivo* activation and differentiation of naïve B cells into functional antibody-secreting cells were confirmed through ELISpot analysis, demonstrating a significant increase in total immunoglobulin (Ig) production after seven days of stimulation in comparison to unstimulated PBMCs (Fig. S7A). In addition, we monitored the immunophenotypic evolution of B cells by flow cytometry (Fig. 3B, Fig. S7B). Across the three donors, an average 94% of live cells were non-activated naïve B cells on day zero (D0), while by day three (D3), over 90% of the naïve B cells were activated. By day seven (D7), the composition of cells included 3.6% non-activated naïve B cells, 39% activated naïve B cells, 4.6% unswitched memory B cells, 9.2% memory B cells, 4.8% plasmablasts, and 0.2% plasma cells. At two weeks (D14), the distribution shifted to 21% memory B cells, 9.6% plasmablasts, and 3.3% plasma cells, with a majority constituting IgD-CD27- exhausted memory cells (Fig. 3C).

**Figure 3.**
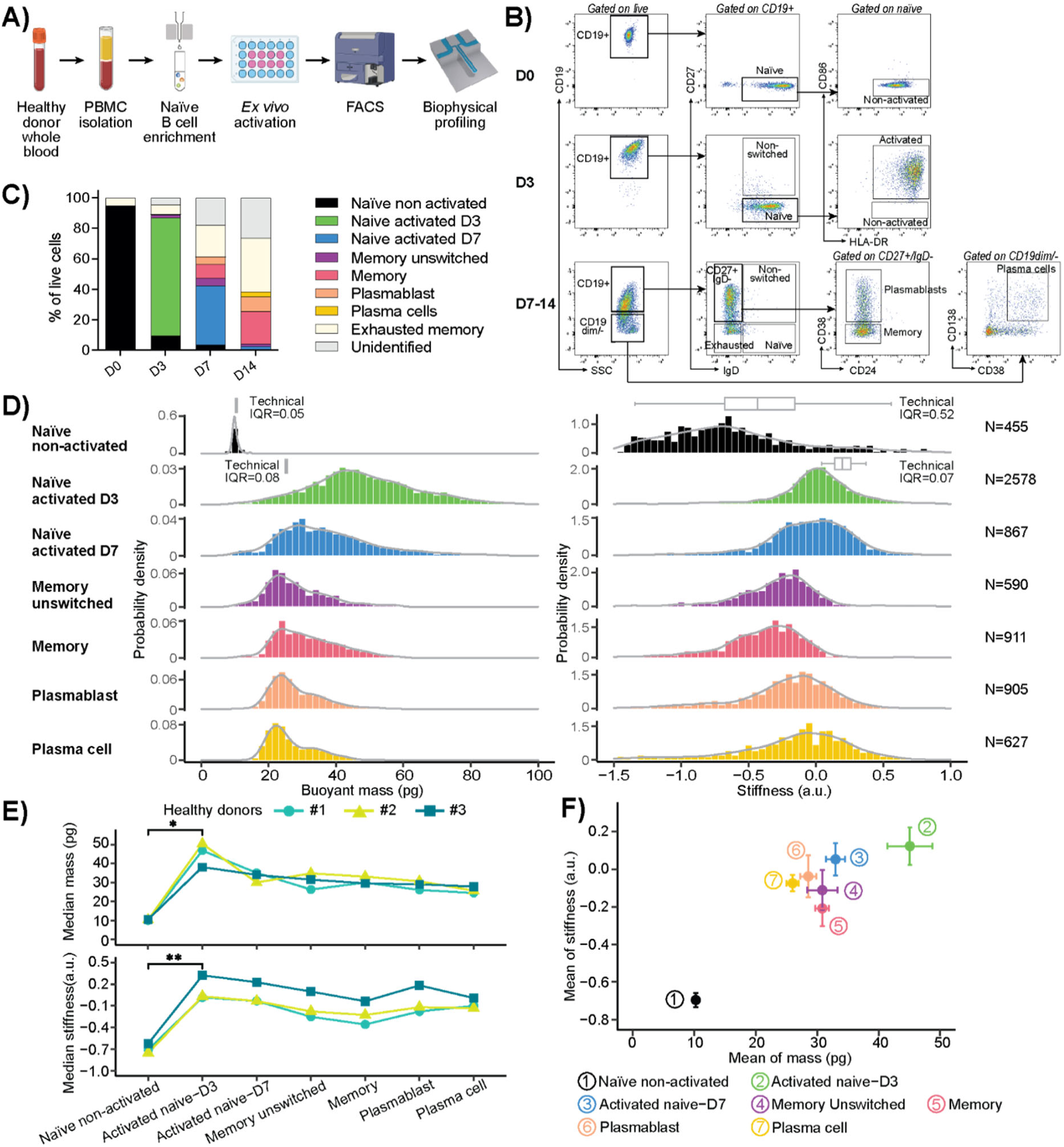
Mature B-cells undergo significant biophysical change after activation. **(A)** Schematic workflow showing the isolation of naïve B cells from PBMCs of three healthy donors, followed by *ex vivo* stimulation for immunophenotyping and biophysical profiling. Created with BioRender. **(B)** Summary of the flow gating strategy used to characterize the different B cell populations. Naïve B cells and stimulated B cells at various differentiation stages were identified by immunophenotyping and sorted by FACS on days 0, 3, 7, and 14. The full panel is shown in Supplemental Fig.2 **(C)** Mean proportion of B cells at different differentiation stages following *ex vivo* stimulation (n=3). **(D)** Representative histogram distributions of single-cell buoyant mass (left) and stiffness (right) of each mature B cell population measured by the SMR, from one experiment out of three biological replicates using cells from 3 healthy donors. The number of single cells measured in each condition is indicated on the right. Technical IQRs were measured by trapping single cells (naïve B cell or L1210 for naïve activated D3) on the SMR (Fig. S3F- G). **(E)** Changes in cells’ buoyant mass (above) and stiffness (below) across three independent *ex vivo* simulations using human PBMCs from three different donors. \**P* < 0.05, \*\**P* < 0.01 as compared between indicated groups (Student’s *t*-test). **(F)** Correlation plot of cells’ buoyant mass and stiffness at each stage of differentiation. Data are presented as mean ± SD of the three biological replicates depicted in (E).

Subsequent biophysical analyses of these distinct stages of differentiation performed using the SMR demonstrated striking changes post-activation. The mass of naïve B cells quadrupled following activation and then decreased by half by the unswitched memory B cells stage (Fig. 3D). Furthermore, mature B cells exhibited an increase in stiffness (*p*-value=0.0077, Fig. 3E). As cells progressed to unswitched memory B cells and memory stages, stiffness decreased but rebounded at the plasma cell phase. Interestingly, while most MCL derive from naïve pre-germinal center B cells (*27*, *47*, *48*), the biophysical profiles of the three PDX models of MCL DFBL-96069, DFBL-39435 and DFBL-91438 were more similar to those of more mature B-cell states. This suggests a reprogramming of cellular mechanisms governing mass and stiffness in MCL cells. The consistency of these biophysical changes was confirmed across three independent *ex vivo* activation and differentiation experiments conducted with PBMCs from three separate donors (Fig. 3E). Additionally, composite biophysical profiles incorporating dual parameters of cell mass and stiffness effectively track B-cell differentiation states (Fig. 3F, Fig. S7C), further supporting the potential for dynamic biophysical characteristics to serve as biomarkers of lymphoid developmental state (*23*, *49*).

### BCR pathway perturbations drive changes in cell mass and stiffness in a MCL cell line

Given the strong correlation between mass, stiffness, and B-cell activation, we sought to define the molecular drivers of phenotypes induced by therapeutic perturbation. To examine the impact of BCR pathway inhibition, we treated Jeko-1 cells, which exhibit high sensitivity to Bruton’s Tyrosine Kinase inhibitors (BTKi)(*50*, *51*), with acalabrutinib, a selective covalent BTKi that blocks BCR signaling. BTK plays a critical role in the BCR pathway and is targeted by inhibitors such as ibrutinib and acalabrutinib in MCL and other B-cell malignancies (*21*, *52*). Treatment of Jeko-1 cells with 0.5 µM of acalabrutinib decreased both mass and stiffness after 24 hours compared to DMSO-treated cells (Fig. 4A, B), while a reduction in stiffness alone was observed after one hour. Conversely, stimulation of Jeko-1 wide-type cells with anti-IgM to activate the BCR signaling pathway resulted in an increase in cell mass after 24 hours, while stiffness increased as early as 10 minutes post-stimulation (Fig. 4C), consistent with our previous findings (*53*). These observations suggest that while cell mass and stiffness are often interrelated, they can be independently modulated by BCR pathway perturbation on different timescales. Cell mass reflects total cellular composition, whereas stiffness primarily reflects membrane composition. Based on their established oncologic relevance and our scSMR-RNA-Seq data (Fig. 2D), we prioritized *BLK* and *CD79A* for further study. BLK and CD79A are critical components of the BCR signaling pathway. CD79A is an essential component of the BCR, and BLK functions as an Src-family kinase, which phosphorylates the immunoreceptor tyrosine-based activation motifs of the transmembrane proteins CD79A and CD79B to mediate signal transduction (*54*, *55*). We engineered the Jeko-1 cell line to overexpress BLK (Jeko-1 +BLK) or CD79A (Jeko-1 +CD79A). Following puromycin selection, the expression of GFP was confirmed by flow cytometry (Fig. S8A) and the overexpression of *BLK* and *CD79A* mRNA and protein was confirmed by RT-qPCR and western blot, contextualizing subsequent biophysical assessments (Fig. 4A). Importantly, neither BLK nor CD79A overexpression significantly affected cell cycle distribution or cell proliferation (Fig. S8B-D). BLK transcript and protein levels were increased four to five fold in Jeko-1 +BLK compared to Jeko-1 GFP-overexpressing controls (Jeko-1 +GFP), while CD79 was increased around two fold in Jeko-1 +CD79A (Fig. 4D, E). BLK or CD79A overexpression led to the activation of key downstream BCR signaling components in comparison to Jeko-1 +GFP, as evidenced by increased BTK and PLCγ2 phosphorylation, with higher relative pBTK/BTK and pPLCγ2/PLCγ2 levels (Fig. S8E). Jeko-1 cells overexpressing BLK or CD79A displayed an increase in both mass and stiffness, in contrast to control cells overexpressing GFP (Fig. 4F). These findings support a mechanistic role for BCR signaling through BLK and CD79A, in governing the biophysical properties of MCL cells, and highlight the ability of BTKi to abrogate their effects.

**Figure 4.**
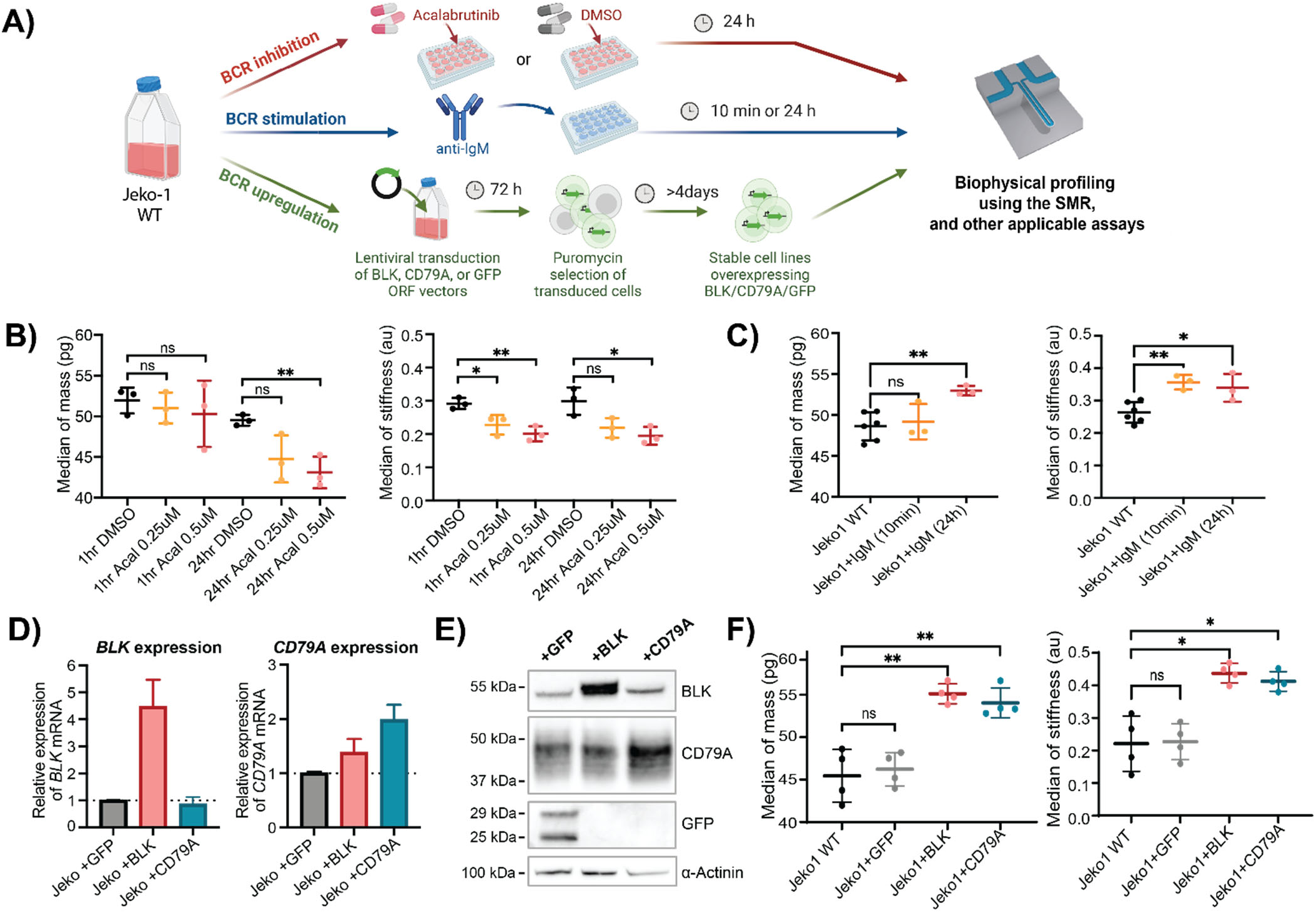
BCR pathway perturbations drive changes in cell mass and stiffness in Jeko-1 cells. **(A)** Schematic illustrating BCR inhibition via acalabrutinib treatment (top panel), BCR stimulation through anti-IgM treatment (middle panel) and BCR pathway enhancement through *BLK* and *CD79A* overexpression (bottom panel) in Jeko-1 wild type (wt) cells for biophysical profiling. BCR inhibition was confirmed by western blot, and stable overexpression in cell lines was validated using RT-PCR, western blot, and flow cytometry. Functional assays (cell cycle and proliferation) further assessed GFP, BLK or CD79A-expressing cells. Created with BioRender. **(B)** Median single- cell mass and stiffness measurements obtained using the SMR from >500 Jeko-1 wild type cells after 1h or 24h of *ex vivo* treatment with DMSO, 0.25 µM, or 0.5 µM acalabrutinib. Data are presented as the median ± SD of three biological replicates. **(C)** Median single-cell mass and stiffness measurements from >500 Jeko-1 wt cells, with or without IgM stimulation for 10min or 24h, assessed using the SMR. Data represent the median ± SD of three biological replicates. **(D)** Relative mRNA expression of *BLK* and *CD79A* measured by RT-qPCR in Jeko-1 overexpressing *GFP*, *BLK*, or *CD79A*. Results are presented as fold change normalized to *GAPDH* mRNA. Each bar represents the mean ± SEM of two independent stable cell lines. **(E)** Representative images of western blot showing BLK, CD79A, GFP and ɑ-actinin protein in Jeko-1 overexpressing *GFP*, *BLK* or *CD79A*. **(F)** Median single- cell mass and stiffness measurements obtained using the SMR from >500 cells of Jeko-1 wild type (WT) and Jeko- 1 cells stably overexpressing GFP, BLK or CD79A. Data are presented as median ± SD of three biological replicates. NS: non-significant, \**P* < 0.05, \*\**P* < 0.01 as compared between indicated groups (Student’s *t*-test).

### Single-cell mass as a biomarker of susceptibility to BTKi in primary patient specimens

Drugs targeting BCR signaling are widely used in MCL therapy, making it a key target for understanding how cellular biophysics can inform treatment strategies. Building upon this premise, we explored the use of these properties as potential *ex vivo* biomarkers of clinical response to BTKi. We analyzed peripheral blood samples from one patient with relapsed/refractory MCL and three treatment- naïve patients (Table S4) participating in an ongoing clinical trial (NCT04855695) assessing the combined effects of acalabrutinib, venetoclax, and obinutuzumab (AVO), which target BTK, BCL2 and CD20, respectively (*56*). Peripheral blood samples were collected before treatment and after one four-week cycle of acalabrutinib monotherapy prior to the addition of venetoclax and obinutuzumab (Fig. 5A). Patients were classified as having either more sensitive or less sensitive disease based on the kinetics of circulating lymphoma response (Fig. S9). PBMCs were isolated from these samples, and MCL cells were subsequently enriched using human CD19-coupled magnetic beads for single-cell mass measurements with the SMR, with immunophenotypic confirmation by flow cytometry (Fig. S10). Notably, for the two patients with more acalabrutinib-sensitive disease (MCL#0491 and MCL#0557), there was a noticeable decrease in the mass of MCL cells during BTK inhibition compared to the pre-treatment state (Fig. 5B). Conversely, for the two patients with less acalabrutinib-sensitive disease (MCL#0528 and MCL#0533), the mass distributions of MCL cells pre- and on-treatment appear to have a similar profile, suggesting that BTK inhibition did not significantly alter the biophysical properties of these MCL cells. Importantly, cell viability remained consistently above 80% across all conditions, and a significant portion of the cells were in G0/G1 phase, indicating the expected state of circulating cells (Fig. S11A-B). This ensures the observed differences in mass are not due to significant cell death or changes in cell cycle distribution.

**Figure 5.**
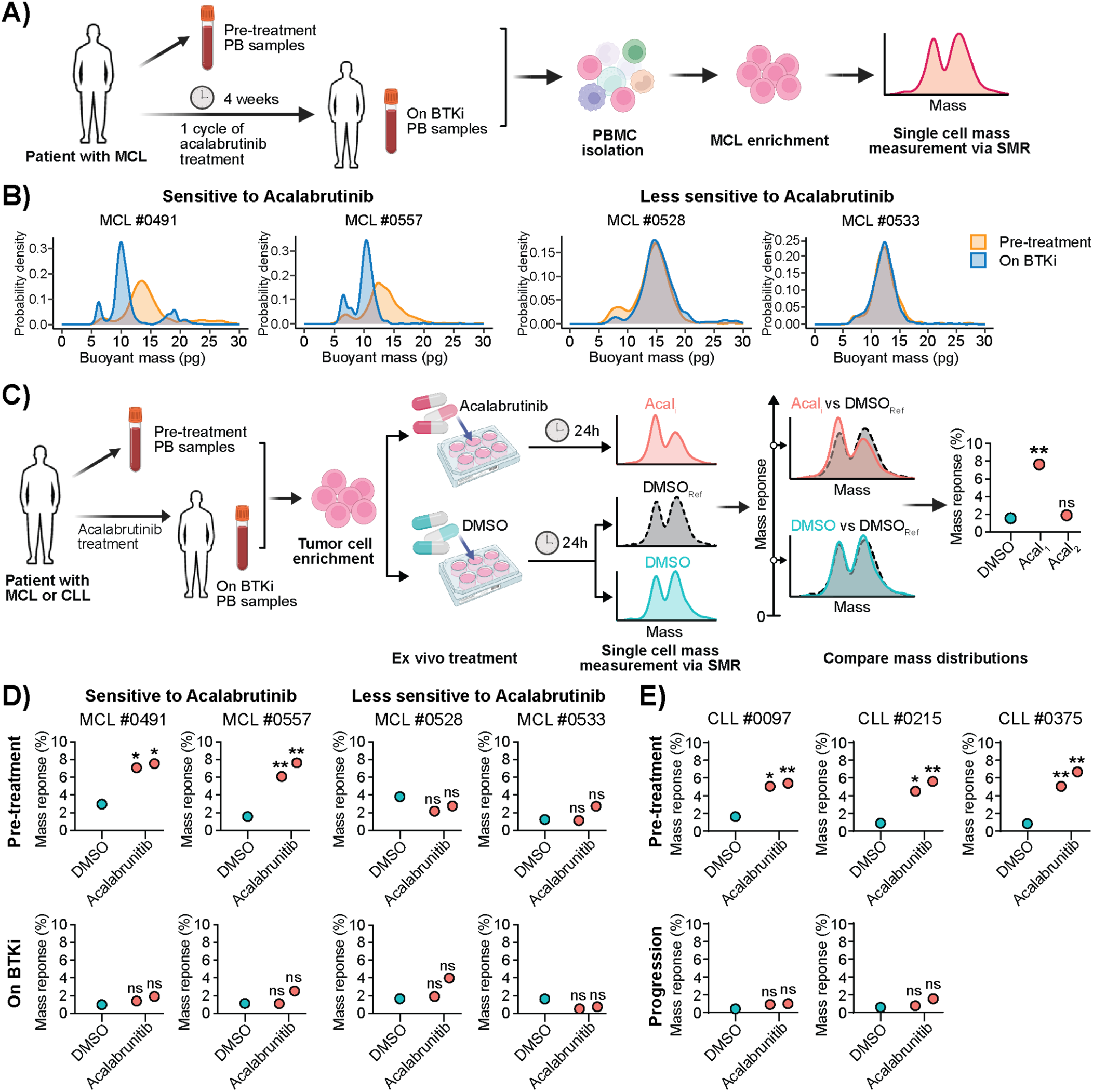
Single-cell mass as a biomarker of susceptibility to pharmacologic inhibition of BCR signaling pathway activity in primary patient specimens. **(A)** Workflow to assess the impact of BTK inhibition on the buoyant mass of MCL cells from patients before and after 4 weeks of acalabrutinib treatment. MCL cells were enriched from PBMC by immunomagnetic depletion. Created with BioRender. **(B)** Single-cell mass distribution of over 1,000 cells measured by the SMR of two acalabrutinib-sensitive MCL primary samples (left) and two less-sensitive MCL primary samples (right) before and after 4 weeks of treatment with the BTKi acalabrutinib as outlined in workflow (A). The samples were categorized as sensitive or less sensitive based on changes in their white blood cell count following BTKi treatment (Fig. S9). **(C)** Schematic workflow to examine the impact of BTK inhibition on the buoyant mass of MCL or CLL cells pre- and post-acalabrutinib *in vivo* treatment, followed by 24h of *ex vivo* drug treatment with DMSO or acalabrutinib. Created with BioRender. One sample of DMSO-treated cells serves as the “reference” distribution. Mass response signals are calculated using EMD analysis, comparing the reference to a second replicate of DMSO-treated cells (“Ref vs DMSO”), and to acalabrutinib-treated cells (“Ref vs Acalabrutinib”) respectively. A *p*-value is determined by comparing the difference between these two mass response signals against a decision threshold. **(D)** Mass response signals of two acalabrutinib-sensitive and two less-sensitive MCL primary samples obtained (bottom), which were subjected to 24h of *ex vivo* drug treatment in duplicate with DMSO or acalabrutinib, as outlined in workflow (C). **(E)** Mass response signals of three CLL primary samples at pre-treatment (top) and progression (bottom) timepoints, followed by 24h of *ex vivo* drug treatment in duplicate with DMSO or acalabrutinib as detailed in workflow (C). NS: non-significant, \**P* < 0.05, \*\**P* < 0.01.

We then exposed the same samples to 0.1 µM acalabrutinib or DMSO in duplicate for 24 hours before repeating the SMR measurement (Fig. 5C). For single-cell mass analysis and comparison, only viable cells were selected based on their viability gating (Fig. S12A). Cell viability remained consistent between DMSO and acalabrutinib treatment (Fig. S12B). For MCL cells derived from the two patients with more acalabrutinib-sensitive disease prior to *in vivo* treatment, there was a notable decrease in mass after 24 hours of *ex vivo* acalabrutinib treatment in contrast to MCL cells derived from less acalabrutinib-sensitive disease (Fig. S13). However, after these patients underwent one cycle of acalabrutinib treatment, subsequent *ex vivo* exposure to the drug for 24 hours did not result in further reduction in mass in either group, suggesting that the *in vivo* treatment had already reached its maximal effect on mass distribution. We assessed the statistical robustness of mass trajectories upon *ex vivo* treatment by comparing the single-cell mass distribution of acalabrutinib-treated cells to that of DMSO-treated cells using “mass response” signals, which quantify the statistical similarity between single-cell mass distributions (Fig. 5C, Methods)(*57*). A small mass response signal is observed when comparing similar distributions, while a larger mass response signal indicates diverging distributions. To account for potential deviations in single-cell mass distributions due to instrument noise, sampling error or phenotypic drift, which may be statistically significant but not biologically meaningful, we measured DMSO-treated cells in duplicate, using one sample as the “reference” distribution to capture these treatment-independent variations. MCL cells obtained prior to *in vivo* treatment from two patients who were more sensitive to acalabrutinib demonstrated significant mass responses after 24 hours of *ex vivo* acalabrutinib exposure (*p*-value =0.0125 & 0.035 in MCL #0491, 0.0025 & 0.005 in MCL#0557). In contrast, mass responses were not significant in persistent MCL cells obtained from patients following a month of *in vivo* treatment or in cells from patients less sensitive to acalabrutinib (Fig. 5D, Fig. S13).

To extend the analysis to another BTKi, the non-covalent inhibitor pirtobrutinib (*58*), we isolated MCL cells from the peripheral blood of a patient with relapsed/refractory MCL pre- and post-4 weeks of pirtobrutinib monotherapy. Single-cell biophysical analysis revealed a reduced mass in MCL cells on pirtobrutinib treatment. This finding aligns with clinical observations, where the patient experienced resolution of neutropenia and symptomatic improvement (Fig. S14A-B). These results raise the possibility that SMR-based mass analysis may be extensible to multiple inhibitors. Intriguingly, such changes in mass distribution were not observed in the non-MCL fraction of cells from the same peripheral blood specimens post-BTK inhibitor treatment, indicating a specific response in the tumor cells.

### Extending the SMR analysis as a predictive biomarker to additional B-cell malignancies

BTKi is standard of care in multiple B-cell malignancies, including chronic lymphocytic leukemia (CLL)(*59–61*). There are few biomarkers to predict treatment response or resistance to BTK inhibition. Therefore, we examined how BTK inhibition affects the mass of peripheral blood CLL cells, adopting a similar experimental approach (Fig. 5C). Primary CLL cells obtained from three patients prior to treatment (Fig. S15A, B) exhibited a significant decrease in mass after 24-hour *ex vivo* exposure to acalabrutinib compared with DMSO controls (Fig. 5E and Fig. S15C; *p*-value<0.05). In contrast to treatment-naïve patients’ samples, CLL cells from patients who had progressed on acalabrutinib treatment exhibited no notable mass alteration following an additional 24-hour *ex vivo* treatment. As with the MCL cells from peripheral blood, the large majority of CLL cells in all samples were in G0/G1, indicating that the changes in mass are related to effects on the targeted signaling pathway rather than non-specific cell cycle effects. These findings suggest that *ex vivo* mass measurements have the potential to predict *in vivo* response to acalabrutinib, offering a potential strategy for identifying patients who may benefit from BTKi.

## DISCUSSION

Our approach links mass and stiffness over the course of *ex vivo* B cell maturation and establishes a dynamic reference of functional biophysical properties across lymphoid lineage development. This complements the recent comprehensive atlas of static cell size and mass measurements across human tissues *in vivo* (*62*). Additionally, the integration of cell mass and stiffness as dual parameters effectively characterizes B cell differentiation states. These data highlight the potential for biophysical features to serve as biomarkers of lymphoid developmental state.

Here we focused on MCL as an archetype of malignant B cell inter- and intra-tumor heterogeneity, to advance the utility of our platform for defining biophysical features and their mechanistic underpinnings. Specifically, we linked mass, stiffness, and gene expression within individual lymphoma cells using scSMR-RNA-Seq to establish associations between biophysical features and oncogenic molecular programs underlying clinically relevant malignant B-cell phenotypes. We show that therapeutic or genetic manipulation of the BCR pathway drives biophysical changes independent of survival and cell cycle, and that treatment of patients with either MCL or CLL results in rapid *ex vivo* changes in malignant cell mass only among responders, supporting ongoing efforts to develop cell mass as a predictive biomarker of clinical response.

Our data demonstrate the ability of single-cell biophysical-transcriptional profiling to identify functionally significant signaling pathways and its potential to reveal or validate therapeutic targets in primary human cancer cells. Notably, genes associated with cell proliferation, division, and cytoskeleton organization showed positive correlations with both mass and stiffness. The unique profile of the PDX model DFBL- 96069, marked by the expression of genes associated with cell cycle arrest and negative regulators of proliferation, may underlie its distinctive single-cell biophysical profile, which was characterized by lower mass and stiffness. The correlation between biophysical properties and key pathways governing B-cell development such as BCR signaling, whose role in MCL tumor biology is well-established, supports the clinical relevance of this approach. We selected the BCR signaling pathway for deeper interrogation because of its effective targeting by standard MCL therapies. Our data demonstrate that the overexpression of *BLK* and *CD79A* is sufficient to increase cell mass and stiffness, corroborating their roles in MCL biology and response to BTKi. Our findings additionally revealed that biophysical properties correlate with cell proliferation and cytoskeleton organization plus other oncogenic signaling pathways such as those governing cell adhesion. Changes in cell adhesion properties can alter tumor cell interactions with their microenvironment, influencing their ability to disseminate or persist during treatment. The observed anti-correlation between *JUN*, *TNF*, and *DUSP1* expression and cell mass suggests these genes might reflect treatment resistance or stress responses. These correlations with biophysical features merit further evaluation in larger cohorts to clarify mechanisms of resistance and persistence in MCL. Future efforts leveraging the non-destructive nature of biophysical profiling with the SMR to capture biological features orthogonal to gene expression, such as chromatin conformation and phosphoproteomics, may create additional unique opportunities to elucidate novel mechanisms of therapeutic response and resistance.

Our findings also highlight the ability of high-precision single-cell biophysical dynamics to provide insights into therapeutic response throughout the treatment continuum. This has several potential clinical applications. First, given the heterogeneity of MCL disease biology, improved baseline disease characterization is needed to risk-stratify patients more accurately and inform treatment selection and intensity. Our data demonstrate that MCL tumor populations from different PDX models exhibit unique cell mass and stiffness profiles. Validating the correlation between specific profiles and clinical responses would create a rationale for incorporating baseline biophysical properties into upfront MCL disease characterization and diagnostic risk stratification models. Second, predicting response to therapy whether at baseline or at relapse would enable precision approaches that tailor therapy to an individual patient’s tumor biology. The distinct biophysical profiles we observed in malignant cells from patients responsive to BTKi suggest that mass could potentially serve as a functional readout of BTKi sensitivity. This was true not only of MCL but also CLL, the most common form of leukemia in adults, comprising nearly a third of all leukemias in North America and Europe. Our results therefore justify a more systematic characterization of biophysical responses to *ex vivo* drug treatment to correlate with *in vivo* sensitivity and clinical outcomes. The SMR technology has previously been used to assess drug responses in primary tumor specimens via single-cell mass measurements (*57*). With its rapid turnaround time (∼24 hours), SMR-based biophysical profiling could be a valuable tool for serial monitoring of therapeutic response and informing response-adapted treatment regimens for MCL patients.

Since persistent tumor cells eventually give rise to clinical relapse, there has been significant interest in interrogating MCL in clinical remission on or after treatment at the point of minimal residual disease (MRD). Understanding the biology of these cells represents an attractive prerequisite for the rational design and selection of novel therapies to suppress or even eradicate MRD. Our approach for biophysical interrogation can be performed using low-input specimens (∼2000 cells)(*63*). It therefore presents a compelling opportunity for characterizing MRD that has yet to be investigated in MCL.

Our results provide an important foundation for further developing clinical applications of biophysical properties, albeit with caveats. Here we analyzed primary samples from a limited number of patients who reflect only a subset of the clinical, genetic, and transcriptomic heterogeneity observed in MCL. Additionally, the current single-cell sorting system following the SMR measurement and the SMART-Seq2 sequencing method, while advantageous for its high-resolution transcriptomic analysis, limits our paired biophysical-transcriptomic analysis to hundreds of cells. To fully capture the clinical and biological heterogeneity of MCL, profiling a larger number of cells may be necessary to adequately capture MCL subclones. Moreover, our study employed *ex vivo* drug treatment assays to assess therapeutic responses. However, it is important to acknowledge that these assays are inherently subject to biological biases. The processes involved in tumor cell dissociation and enrichment can alter the native tumor microenvironment and impact cell behavior and drug response. This limitation underscores the necessity of interpreting our findings with caution and highlights the importance of developing more representative models that preserve the complexities of the tumor microenvironment.

Finally, we showed promising data that assessment of biophysical properties, particularly single-cell mass, could provide a rapid readout of BTKi sensitivity. While these findings are encouraging, validation in larger patient cohorts will be necessary to establish the potential of single-cell mass measurements as an *ex vivo* surrogate for *in vivo* drug sensitivity. Moreover, given the narrow focus of our study, additional investigation is needed to define the correlation between cell mass and BCR signaling pathway activity, as well as to explore the broader utility of biophysical properties in other diseases or treatment contexts. Therefore, ongoing investigation aims first to validate the correlation between biophysical properties and clinical outcomes systematically in clinical trial cohorts, with dedicated assessment of relapsed/refractory and treatment-naïve MCL, as we are doing in NCT04855695. Such clinical validation would justify statistically powered evaluation of single-cell mass and stiffness as biomarkers of clinical response to regimens involving similar therapies within other B-cell neoplasms (e.g., CLL, diffuse large B-cell lymphoma, follicular lymphoma, marginal zone lymphoma, lymphoplasmacytic lymphoma). Subsequent efforts would then seek to define the ability of serial single-cell biophysical profiling with SMR to predict outcomes and reveal candidate therapeutic targets even in low tumor burden settings including on- or post-treatment residual disease.

## MATERIAL AND METHODS

### PDX generation

All *in vivo* experiments were conducted under Dana-Farber Cancer Institute Animal Care and Use Committee protocol #13-034. A full description of each PDX model is available online at the Public Repository of Xenografts (https://www.cbioportal.org/, CPDM Pan-Cancer Patient Derived Models)(*25*), including clinical history and genomic data. All mice were maintained under a 12-hour light/12-hour dark cycle at constant temperature (23°C), with free access to food and water. For PDX models, viably frozen PDX cells were thawed and washed in phosphate buffered saline (PBS) before tail-vein injection at 1×10^6^ cells per mouse. Female Nod.Cg-*Prkdc^scid^IL2rg^tm1Wjl^*/SzJ (NSG) mice aged 6- to 8-week (Jackson labs, #005557) were used as recipients. Tumor burden was monitored periodically by flow cytometry of peripheral blood. Blood was processed with ammonium–chloride–potassium (ACK, Life Technologies, #A1049201) before staining with antibodies against human CD45 (BD Biosciences, #566026) and human CD19 (Thermo Fisher Scientific, #BDB564456) in PBS with 1 mM EDTA plus 2% fetal bovine serum (FBS). Data were acquired on a BD Fortessa flow cytometer and analyzed with FlowJo Software. Mice were monitored daily for clinical signs of disease and humanely euthanized when they reached a clinical end point. Cells from the spleen, liver, peripheral blood, and bone marrow were collected, ground through a 70 µm filter, and subjected to ACK lysis. Human cells were then selected using the EasySep Mouse/Human Chimera Isolation Kit (Stem Cell Technologies, #19849) according to the manufacturer’s protocol. Human tumor cell number and viability were assessed with Trypan blue staining.

### Cell culture

Jeko-1 and Rec1 cells were cultured in RPMI-1640 (Gibco, #11875-093) supplemented with 20% FBS (Sigma Aldrich, #F2442) and 1% penicillin/streptomycin (P/S). Maver-1 and Mino cells were cultured in RPMI 1640 supplemented with 10% FBS and 1% P/S. Granta-519 cells were cultured in DMEM (Gibco, #11995-065) supplemented with 10% FBS and 1% P/S, and Z-138 cells were cultured in IMEM (Gibco, #12440-053) supplemented with 10% FBS and 1% P/S. HEK293T cells were cultured in DMEM supplemented with 10% FBS, and 1% P/S. The cells were maintained at 37°C under 5% CO_2_. Cell lines were routinely tested for mycoplasma (InvivoGen, #rep-mysnc-50). All the cell lines were purchased from ATCC.

### Primary cells

All studies involving primary patient samples were approved by the Dana-Farber/Harvard Cancer Center institutional review board. Informed consent was obtained in accordance with the Declaration of Helsinki. MCL serial primary blood specimens were obtained from appropriately consented patients treated on a phase 1/2 study, investigator-initiated, single-arm clinical trial (NCT04855695), to assess the safety and efficacy of the combination of acalabrutinib, venetoclax, and obinutuzumab (AVO) in relapsed/refractory (R/R) and untreated MCL. These patients cross-consented to Dana-Farber Cancer Institute tissue banking protocol #21-040. The samples used in our study include peripheral blood obtained at screening and on days one of cycle two, before the start of the obinutuzumab treatment. Peripheral bloods specimens were collected into EDTA vacutainer tubes prior to peripheral blood mononuclear cell (PBMC) isolation by Ficoll-Paque Plus (Thermo Fisher Scientific, #45001749) gradient centrifugation per standard protocols and viably frozen. CLL specimens were generously gifted by Jennifer Brown laboratory at Dana-Farber Cancer Institute. The patients cross-consented to Dana-Farber Cancer Institute CLL tissue banking under an institutional review board approved protocol. PBMCs were isolated per standard protocols and viably frozen. CLL or MCL tumor cells were enriched using EasySep™ Human B Cell Enrichment Kit II Without CD43 Depletion (Stem Cell technology, Catalog #17963) according to the manufacturer’s protocol. Tumor cells were >95% of cells as determined by flow cytometry.

### Biophysical measurement using the SMR

Single-cell mass and stiffness measurements were conducted using the SMR as detailed in Burg et al. (2007)(*52*) and Kang et al. (2019)(*24*). The SMR comprises a cantilever-based mass sensor that also detects acoustic scattering from cells, which is incorporated in an integrated microfluidic channel. As cells traverse this channel, their passage alters the cantilever’s resonance frequency, the changes of which were used to quantify the mass and acoustic scattering from the cells. A parallel volume measurement using coulter counter (Beckman Coulter) was carried out to quantify average cell volume and this was used together with the single-cell mass measurements to carry out cell size normalization of the acoustic scattering signal to calculate cell stiffness, as reported before(*24*). Comprehensive procedural details are available in the cited works.

In preparation for measurements, the SMR system was subjected to a rigorous cleaning protocol to eliminate residual biological debris. Initially, the device was treated with 0.05% Trypsin-EDTA (Invitrogen, #25300054) for 20 minutes, followed by a 3-minute exposure to 5% bleach. Subsequently, the system was rinsed for 5 minutes with deionized water. After cleaning, the channel was passivated using 1 mg/mL of PLL-g-PEG (Nanosoft Polymers, #11354) in water for 10 minutes at room temperature, and then rinsed again for 5 minutes with flow cytometry (FACS) staining buffer (Rockland, #MB-086-0500). Measurements were performed in FACS buffer at room temperature, each lasting no longer than 30 minutes. Between samples, the SMR was briefly flushed with FACS buffer to maintain cleanliness. During the experimental runs, samples were introduced into the SMR via 0.005-inch-inner-diameter fluorinated ethylene propylene (FEP) tubing (Idex, #1576L). Fluid flow within the SMR was regulated using three electronic pressure regulators (MPV1, Proportion Air) and three solenoid valves (SMC, #S070), ensuring consistent differential pressure to maintain uniform shear and consistent data acquisition rates during cell measurement. All system components, including the regulators, valves, and data collection, were controlled via a custom LabVIEW 2020 (National Instruments) software interface. When the cell viability was lower than 85%, the live cells were enriched by negative selection using the dead cell removal kit (Miltenyi Biotec, #130-090-101) and MS separation columns (Miltenyi Biotec, # 130-042-201) prior to the SMR measurement.

### Single-cell density measurement

Cell density and volume were measured using fluorescent exclusion techniques as previously described (*29*). A fluorescent microscope was positioned at the entry to the SMR cantilever to couple single-cell mass and volume measurements. The fluorescence level emitted from the detection region on the SMR was continuously monitored by a photomultiplier tube (PMT, Hamamatsu, H10722-20). Cells were suspended in PBS with 2%FBS and 5mg/mL FITC-conjugated dextran (Sigma, FD2000S-250MG). When there was no cell present at the detection region, the PMT detected a high fluorescence baseline from the fluorescence buffer. When a cell passed through, the fluorescence signal detected by the PMT decreased proportionally to the volume of the cell. Immediately after volume measurement, each single cell flowed through the SMR cantilever, and the corresponding mass was resolved. Single cell density 𝜌 is then computed using the equation as below,

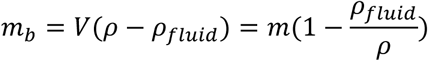

where 𝑉 is the volume of the cell, 𝑚 is the mass and 𝜌 is the density of the cell immersed in a fluid of density 𝜌_fluid_. The fluid density of PBS with 2%FBS and 5mg/mL FITC-conjugated dextran is 1.005 𝑔⋅𝑐𝑚^-3^.

### Histology

Mice tissues were fixed in 10% neutral buffered formalin for 24 hours and then kept in 70% ethanol. Hematoxylin & Eosin (H&E) and immunohistochemistry (IHC) were performed at the Brigham and Women’s Hospital Pathology Core at Harvard Medical School. IHC were performed using anti-human- p53 antibody (clone E26, Abcam, #ab32389), anti-human CD20 antibody (clone E7B7T, Cell Signaling Technology, #48750), anti-mouse/human Ki67 (clone SP6, Biocare, #PRM325) and anti-mouse/human Cyclin D1 (clone SP4, Neomarkers, #RM-9104-S).

### Single-cell RNA-sequencing

Suspension of peripheral blood, bone marrow, liver, and spleen cells isolated from DFBL-96069, DFBL- 39435, and DFBL-91438 were sorted into 96-well PCR plates containing 2 μl lysis buffer, spun down and frozen at −80°C. To generate Smart-seq2 libraries, priming buffer mix containing dNTPs and oligo-dT primers was added to the cell lysate and denatured at 72°C. cDNA synthesis, pre-amplification of cDNA and tagmentation was performed as described previously (*30*, *31*). Representative cDNA from single cells was assessed with an Agilent High Sensitivity DNA kit (Agilent Technologies, #5067-4626). Single-cell cDNA was tagmented and pooled to generate libraries by using an Illumina Nextera XT DNA sample-preparation kit (Illumina, #FC-131-1096) with 96 dual-barcoded indices (Illumina, FC-131-1002). The library cleanup and sample pooling were performed with AMPure XP beads (Agencourt Biosciences, #A63880). Barcoded libraries were purified and quantified using Qubit 4.0 Fluorometer (Invitrogen) according to the Qubit dsDNA HS Assay Kit (Life Technologies, #Q32854), and pooled at equimolar ratios.

Sequencing reads were aligned to the human genome (hg19) using STAR version 2.7.9a with default parameters. Sorted and indexed bam files were passed to htseq-count v2.0.1 for transcript quantification with mode "intersection-nonempty", with supplementary and secondary alignments ignored. Cells with fewer than 1000 total reads or greater than 20% of reads corresponding to mitochondrial genes were removed. Principle component analysis (PCA), nearest neighbor finding, and 2D t-SNE plots were generated using Seurat, and number of genes detected, alignment rate, total library size and percent mitochondrial reads were visualized to identify clusters of low-quality cells. These clusters were removed and the same process applied iteratively until no low-quality clusters remained. Correlation of expression with biophysical measurements was performed using Spearman correlation on the log-normalized counts. Genes were nominated for ontology analysis by converting correlation coefficients to z-scores and selecting all genes with a z-score greater than 2.5 for both mass and stiffness gene lists (n=113 for mass, n=61 for stiffness). Gene ontology enrichment was performed on the top positively correlated genes for each biophysical feature against the Biological Process ("BP") group of ontology gene sets in the "C5" category of the MSigDB database using the enrichGO function of the clusterProfiler package. Multiple hypothesis correction was done using the Benjamini-Hochberg method, and results were ranked by -log10(qvalue). Combined ranking of gene correlation for both mass and stiffness was done by converting the rho coefficients of each gene to z-scores and ranking by the average z-score across biophysical correlates. Empirical confidence intervals for Fig. 2D were determined by randomly pairing gene expression profiles and biophysical profiles across 10,000 iterations and taking the 95th percentile correlation coefficients per gene.

### Bulk RNA-sequencing

RNA was extracted using the RNeasy Plus Mini Kit (Qiagen, #7413) according to the manufacturer’s protocol. Total RNA quality was checked using the Bioanalyzer with the Agilent RNA 6000 Pico Kit (Agilent, #5067-1535). The mRNA library was prepared using poly A enrichment and sequenced using an Illumina NovaSeq PE150 system at the Novogene Bioinformatics Technology Co. Ltd, Sacramento CA 95817.

Sequencing reads were aligned to the human genome (hg19) using STAR version 2.7.9a using default parameters, and transcripts were quantified using htseq in the same manner as the single-cell dataset. Samples were checked for library complexity, alignment rate, number of genes detected and percent mitochondrial reads, but no samples needed to be removed due to quality. Transcriptomes were normalized using DESeq2 with default parameters, and a variance stabilizing transformation was applied to the data, and the transformed and normalized counts were used for downstream analysis.

### Flow cytometry

The cells were stained in Brilliant Stain Buffer (BD Biosciences, # 566349) with the diluted antibodies listed in Table S5 to their predetermined optimal concentrations. The cells were then washed twice with PBS supplemented with 2% FBS and 0.2% EDTA. Stained cells were analyzed on a BD LSR Fortessa flow cytometer using BD FACSDiva software (Dana-Farber Cancer Institute Flow Cytometry core), and data were analyzed using FlowJo v10.

### B cell activation

Apheresis leukoreduction collars from anonymous healthy platelet donors were obtained from the Brigham and Women’s Hospital Specimen Bank under an institutional review board–exempt protocol. PBMCs were isolated with Ficoll-Paque Plus (Thermo Fisher Scientific, #45001749) using the manufacturer’s recommended protocol. The PBMC layer was isolated, subjected to ACK lysis (Life Technologies, #A1049201), and washed with PBS. Naïve B cells were isolated using EasySep™ Human Naïve B Cell Isolation Kit (Stem Cell, #17254) according to the manufacturer’s protocol. The naïve B cells were seeded at 1x10^5^ to 2.5x10^5^/ml in a 6-well plate and cultivated in the ImmunoCult™ Human B Cell Expansion Kit (Stem Cell, #100-0645) for 14 days according to the manufacturer’s protocol. The density was adjusted to 1x10^5^/ml every 2 to 4 days. The immunophenotype was confirmed by flow cytometry after 0, 3, 7, and 14 days of cell culture. The viability was assessed by Trypan blue, and the production of total IgG at day 7 was measured using Human IgG ELIspot Basic (Mabtech, Inc. # 3850-2A). On days 7 and 10 to 14, the cells were sorted on a FACSAria II SORP fluorescence-activated cell sorter (BD Biosciences) (Table S5). Naïve B cells were enriched using CD19+/IgD+/CD27-/HLA-DR+/CD86-,activated naïve B cells using CD19+/IgD+/CD27-/ HLA-DR+/CD86+, the memory unswitched B cells using CD19+/IgD+/CD27+, the memory B cells using CD19+/IgD-/CD27+/CD38-, the plasmablasts using CD19+/IgD-/CD27+/CD38+, and the plasma cells on CD19dim or CD19-/CD38+/CD138+. The cells were then washed and resuspended in PBS supplemented with 2% FBS and profiled using the SMR.

### ELISpot assays

The ELISpot assay was performed using the Human IgG ELIspot Basic kit (Mabtech, Inc. #3850-2A) according to the manufacturer’s protocol. Briefly, the coating monoclonal antibodies MT91/145 were incubated on the ELISpot plate overnight for total IgG detection. The plate was washed and blocked, and unstimulated PBMC or stimulated naïve B cells were then washed, resuspended in RPMI +10% FBS, and incubated on the coated plate at 37°C in a humidified incubator with 5% CO2 for 24 hours. The wells were then washed 5 times with PBS, incubated with detection antibodies and Streptavidin-ALP, and developed with BCIP/NBT substrate solution until spots appeared.

### BCR pathway stimulation with anti-IgM

Jeko-1 wild-type cells were obtained from the cell culture incubator and treated with 5 μg/mL anti-IgM (Southern Biotech, Cat. No 2022-01) in an Eppendorf tube at 37°C for 10 mins. Cells were then immediately loaded into the SMR for measurement. After measurement of the stimulated Jeko-1 cells, a new batch of the same cells was measured on the SMR without IgM treatment. The SMR was briefly washed with PBS between each experiment.

For 24-hour stimulation, Jeko-1 wild-type cells (1 million per mL) were incubated with 5 μg/mL anti-IgM for 24 hours at 37°C under 5% CO_2_. Following treatment, cells were harvested and subjected to single-cell mass and stiffness measurements using the SMR, as previously described. Cell viability was assessed using Trypan blue staining.

### Generation of overexpression cell lines

To produce Jeko-1 stable cell lines overexpressing GFP, BLK or CD79A, lentiviral particles expressing a CMV promoter, the respective cDNA and a puromycin resistance gene were purchased from Genecopoeia. Lentivirus were produced by co-transfecting HEK-293T cells with HIV packaging mix (GeneCopoeia, #LT001) and the expression vector EX-A0751-Lv241 containing the open reading frame (ORF) expression clone for *BLK* (NM_001715.2), EX-G0034-Lv241 containing the ORF for *CD79A* (NM_001783.3) or the pReceiver EX-EGFP-Lv241 containing *EGFP* cDNA, using EndoFectin lenti transfection reagent according to the manufacturer instruction (GeneCopoeia, #LT001). After 48 hours, the viral particle-containing supernatant was harvested, filtered through a 0.45 μm protein filter and concentrated using lenti-X™ Concentrator (Takara Bio, #631231) according to the manufacturer’s instructions. The lentiviruses were then added to Jeko-1 cells at different titers, together with 8 μg/mL polybrene (Santa Cruz Biotechnology, #SC-134220). Jeko-1 and transduced cells were selected 72 hours after in their respective complete medium supplemented with 2 ug/ml puromycin (Life Technologies, #J67236.8EQ). The expression of GFP in cells transduced with EX-EGFP-Lv241 was assessed by flow cytometry.

### Reverse Transcription quantitative Polymerase Chain Reaction (RT-qPCR)

Total RNA was extracted from cells using RNeasy Plus Mini Kit (Qiagen, # 74134) as per the manufacturer’s instruction. Then, 500 ng of RNA was transcribed into cDNA using the iScript RT Supermix (Bio-Rad Laboratories, #1708840). qPCR reactions were carried out using the following primers: BLK 5’-CACCGGAGAAGAATTCATCTGGGAC-3’ (forward) and 5’- AAACGTCCCAGATGAATTCTTCTCC-3’ (reverse), CD79A 5’-CACCGCCACTGGGAGAAGATGCCTG- 3’ (forward) and 5’-AAACCAGGCATCTTCTCCCAGTGGC-3’ (reverse), GAPDH 5’-ACCCACTCCTCCACCTTTGA-3’ (forward) and 5’-CATACCAGGAAATGAGCTTGACAA-3’ (reverse).

Quantitative real-time PCR was performed on a CFX96 Real-Time system (Bio-Rad) using PowerUp SYBR Green Master Mix (Life Technologies, #A25742). Target gene expression was normalized to the mean Ct values of the housekeeping gene GAPDH. All values were then normalized to the control sample.

### Western blotting

Jeko-1 overexpressing cells were lysed in RIPA buffer (Sigma Aldrich, # R0278) supplemented with anti-protease and anti-phosphatase cocktails (Cell Signaling Technology, #5872S) to obtain whole cell lysates. Cells were lysed on ice for 30 minutes, and spun down at 4°C 13,000rpm for 10 minutes. Protein concentration was determined using Bradford Protein Assay Kit (Thermo Fisher Scientific, #23225). Equal amounts of protein were boiled in 1X SDS loading dye (Bio-Rad) at 90°C for 10 minutes prior to loading into 4-12% Bis-Tris polyacrylamide gels mini protein gels (Thermo Fisher Scientific, #NP0323BOX). Proteins were then transferred on nitrocellulose membrane using iBlot 2 transfer stacks (Life Technologies, #IB23001). Membranes were blocked 1 hour at room temperature using pierce Clear Milk Blocking Buffer (Life Technologies, # 37587) or 5% bovine serum albumin (BSA, Cell Signaling Technology, #9998S) in TBS-tween, and incubated overnight at 4°C with gentle agitation with the following primary antibodies: anti-BLK (E8T1B) antibody (1:1,000, Cell Signaling Technology, #66002S), anti-CD79A (D1X5C) XP® (1:1000, Cell Signaling Technology, #13333S), anti-GAPDH (D16H11) XP® (1:1,000, Cell Signaling Technology, #5174S), anti-alpha-Actinin antibody (1:1,000, Cell Signaling Technology, #3134S), anti-GFP (1:1,000, Abcam, #ab290), anti-Btk (D3H5) (1:1000, Cell Signaling Technology, #8547S), anti-phospho-Btk (Tyr551) (E5Y6N) (1:2,000, Cell Signaling Technology, #18805), anti-PLC-gamma-2 (Cell Signaling Technology, #3872S) and anti-Phospho-PLC-gamma-2 (Tyr1217) (Cell Signaling Technology, #3871S). Membranes were washed with TBST and incubated with HRP-conjugated anti-rabbit (1:3,000, Bio-Rad, #1706515) supplemented with 5% milk (LabScientific, #M0841) or BSA. Signaling was detected by Western ECL Substrate (Life Technologies, #32109) with ChemiDoc MP Imaging System (Bio-Rad) or ImageQuant 800 (Cytiva).

To evaluate downstream BCR signaling pathways, additional western blot analyses were conducted following the same procedure, with the exception that, when specified, Jeko1 cells were pre-treated with Lipopolysaccharides (LPS) (Sigma Aldrich, #L2630-10MG) at a final concentration of 1 µg/ml in RPMI medium supplemented with 20% FBS and 1% P/S for 24 hours before transduction. On the day of transduction, the LPS solution was removed, and the cells were counted and assessed for viability. The cells were then assessed by RT-qPCR, Western blot and SMR (Fig. S8E-F).

### BTKi *ex vivo* drug treatment

To assess the impact of BTK inhibition on cellular biophysical properties, Jeko-1 wild-type cells were treated with 0.25 µM or 0.5 µM acalabrutinib (Selleck, # S8116) or an equivalent volume of DMSO (Sigma Aldrich, #D8418) and incubated for 24 hours at 37°C under 5% CO2. Following treatment, cells were harvested and subjected to single-cell mass and stiffness measurements using the SMR, as previously described.

For primary specimens, after isolation of PBMC, CLL or MCL tumor cells were enriched using EasySep™ Human B-Cell Enrichment Kit II Without CD43 Depletion (Stem Cell technology, Catalog #17963) according to the manufacturer’s protocol. The immunophenotype of the MCL or CLL cells was confirmed using flow cytometry. MCL cells were plated in duplicate at a concentration of 1 million cells per mL, treated with 0.1 µM acalabrutinib or DMSO and incubated for 24 hours at 37°C under 5% CO_2_. CLL cells were plated in duplicate, when possible, at a concentration of 1 million cells per mL, treated with 2 µM acalabrutinib or DMSO and incubated for 24h at 37°C under 5% CO_2_. The cell viability before and after the ex-vivo treatment was assessed by Trypan blue. In parallel, cell lysates were collected for Western blot analysis to confirm BTK pathway inhibition.

### Integrated biophysical and viability assay

To access cell viability, cells were stained using the Zombie Red^TM^ Fixable Viability kit (BioLegend, #423109) and analyzed with the SMR coupled with a fluorescence microscope (fxSMR) previously developed in our lab (*29*). After *ex vivo* treatment, MCL tumor cells were centrifuged at 500 g for 5 minutes and resuspended in 100µL of PBS buffer (without Tris). Zombie Red dye was added to the cell suspension at a ratio of 1 µL per million cells (e.g., 0.5 µL of Zombie Red dye was added to 500,000 cells in 100 µL of PBS). The cells were incubated at room temperature for 15 minutes, then centrifuged at 500 g for 5 minutes and resuspended in 1 mL of PBS supplemented with 2% FBS for single-cell mass measurement. Fluorescence from Zombie Red staining was detected for individual cells using a fluorescence microscope positioned at the entry to the SMR cantilever, allowing simultaneous detection of the single-cell viability and mass.

### Cell cycle analysis and proliferation assays

To assess cell cycle distribution, cells were fixed in 70% ethanol for 24 hours, washed with PBS, and incubated with 300 μl of FxCycle PI/RNase Staining Solution (Life Technologies, #F10797). Flow cytometry was performed on a BD LSR Fortessa flow cytometer using BD FACSDiva software (Dana-Farber Cancer Institute Flow Cytometry core), and analyses were performed using FlowJo software v10. G2/M cell cycle arrest was induced by treating cells with 40 ng/mL nocodazole (Sigma Aldrich, #M1404) for 20 hours, blocking progression at the G2/M phase. Post-treatment, cells were simultaneously stained with FxCycle PI/RNase staining solution and analyzed by flow cytometry for cell cycle distribution and by the SMR for single-cell mass measurements.

To assess cell proliferation, 50,000 cells were plated in triplicate in 96-well plates for 5 days. On day 3, cells were diluted at 1:3, and this dilution was factored into the cell numbers for viability assays. For apoptosis analysis, cells were stained with 1 µg/mL propidium iodide (PI) solution (Life Technologies, #BMS500PI). Cell numbers and PI^+^ population were detected by flow cytometry. Proliferation assays were performed three times following the puromycin selection of the overexpressing cells.

### Statistics

Figure legends indicate specific statistical analyses used. *P*-values were considered statistically significant as follows: \**p* < 0.05, \*\**p*<0.01, \*\*\**p* < 0.001 and \*\*\*\**p* < 0.0001. Results are expressed as the mean ± s.e.m (as noted in the figure legends) of at least three independent experiments. The biophysical distribution analysis in Fig. 5D, E was performed using the Earth Mover’s Distance (EMD) analysis, which quantifies the statistical similarity between two distributions as a “mass response” signal^41^. The mass response signal is a normalized, dimensionless metric calculated using the following equation,

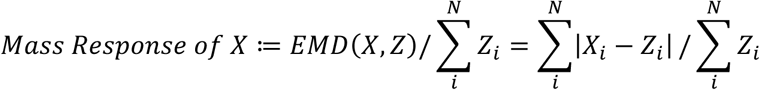

where X and Z are single-cell mass measurements of drug- and DMSO-treated reference cells respectively. N is the number of cells measured in each distribution. Since the number of cells measured (N) to represent each cell population is on the order of thousands, small deviations in the mass distributions due to sampling error, instrument noise, or phenotypic drift can turn out to be statistically significant. Such deviations, however, can be statistically significant without representing a biologically meaningful signal. To circumvent this problem, we define the test statistic, θ, to be the difference between two mass response signals with the following equation,

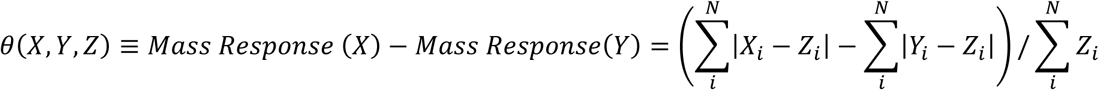

where Y is the single-cell mass measurements of the DMSO-treated control cells. X and Z are defined as above. We compare the test statistic θ to a limit of decision threshold using the bootstrap-t method^41^. All calculations and statistical analyses were carried out using GraphPad Prism (versions 9 or 10) or R (version 4.1.0).

## Acknowledgments

The authors would like to thank all patients who have donated samples for this study; Dr. Jennifer Brown’s laboratory and Stacey Fernandes from the Dana-Farber Cancer Institute for providing CLL specimens.

## Author contributions

Conceptualization: Y.Z., L.D., A.L., D.M.W., S.R.M., M.A.M., Methodology: Y.Z., L.D., A.L., A.I.K., C.E.R., D.M.W., S.R.M., M.A.M., Investigation: Y.Z., L.D., A.L., J.W., I.V., T.P.M., M.Z., E.S., H.L., S.M.D., L.H., J.Z., S.B., R.A.R., M.A., Visualization: Y.Z., L.D., A.L., Data acquisition and analysis: Y.Z., L.D., A.L., J.W., I.V., E.S., H.L., S.M.D., L.H., J.Z., S.B., R.A.R., M.A., Supervision: M.A., M.S.D., A.I.K., C.E.R., D.M.W., S.R.M., M.A.M., Writing—original draft: Y.Z., L.D., C.E.R., S.R.M., M.A.M., Writing—review & editing: Y.Z., L.D., A.L., I.V., T.P.M., E.S., C.E.R., D.M.W., S.R.M., M.A.M., Funding acquisition: A.I.K., D.M.W., S.R.M., M.A.M.

## Supplemental Data & Tables Summary

Supplementary Figs. S1 to S15 Tables S1 to S5

References

