## Supplementary figures and tables for "Integrating Single-Cell Biophysical and Transcriptomic Features to Resolve Functional Heterogeneity in Mantle Cell Lymphoma"

**This PDF file includes:**

Figs. S1 to S15
Tables S4 and S5
Legend for tables S1, S2 and S3
References

**Other Supplementary Materials for this manuscript include the following:**

Tables S1, S2 and S3

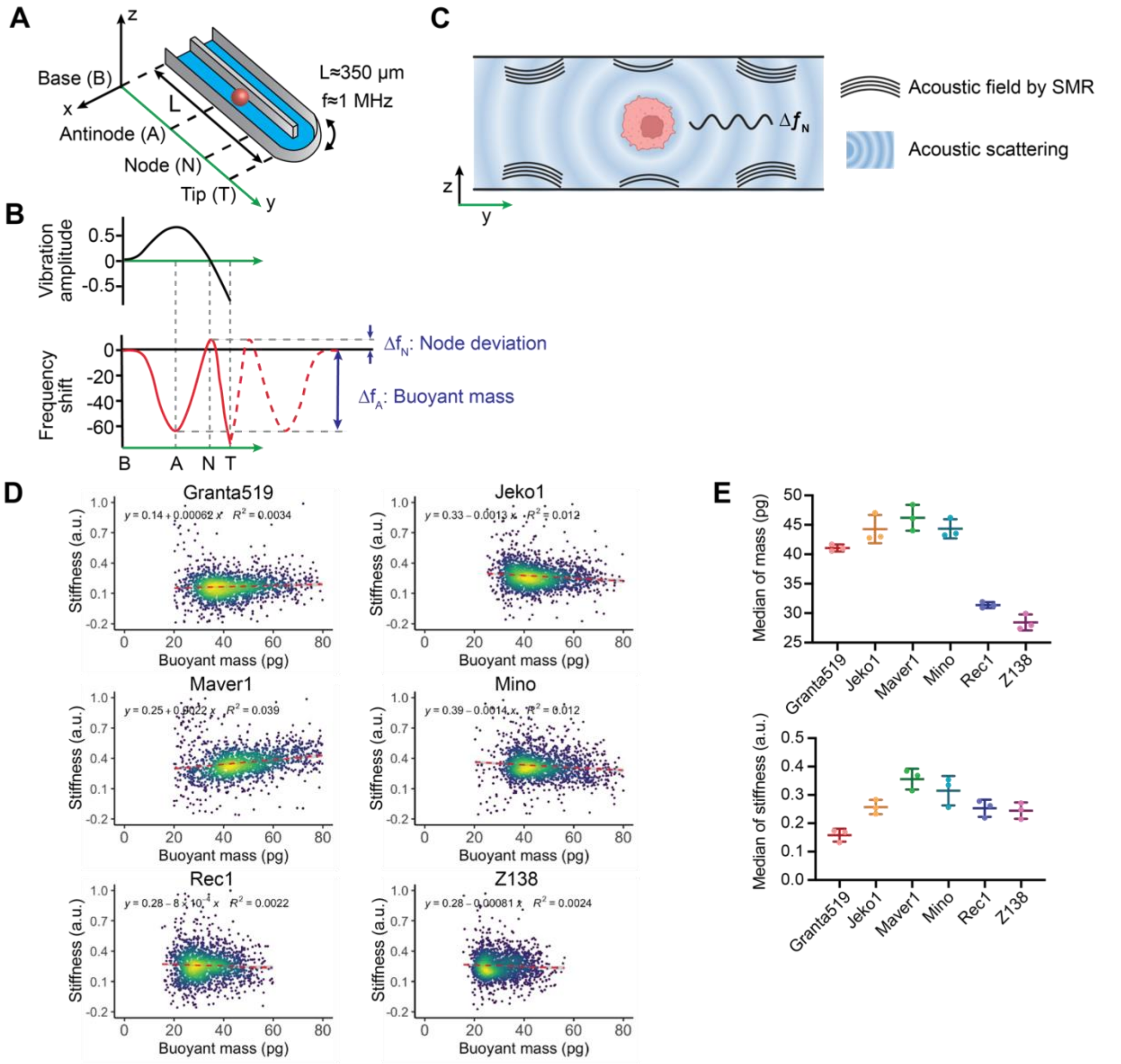


**Fig. S1. Single-cell biophysical measurements utilizing the SMR. A)** Schematic of the SMR with a cell flowing through the embedded fluid channel along the cantilever, which vibrates at resonant frequency f. **B)** Top, normalized vibration amplitude at the second mode. Bottom, resonant frequency shift when a single cell flows along the cantilever in the SMR. The vertical dashed lines indicate the positions of the cell along the cantilever, as in the schematic shown in (A). Buoyant mass is measured at the antinode (A), and node deviation is measured at the node (N). **C)** Illustration of frequency shift due to acoustic scattering. When a cell flowing in the fluid channel interacts with acoustic fields (black waves) generated by the cantilever vibration at resonant frequency, the particle-fluid interaction causes acoustic scatterings (blue waves), which shifts the measured resonant frequency. **D)** Cell mass vs stiffness profiles of six MCL cell lines measured on the SMR. **E)** Median of single-cell buoyant mass and stiffness measurements from >500 cells in each MCL cell line. Data are presented as mean ± SD of three biological replicates.

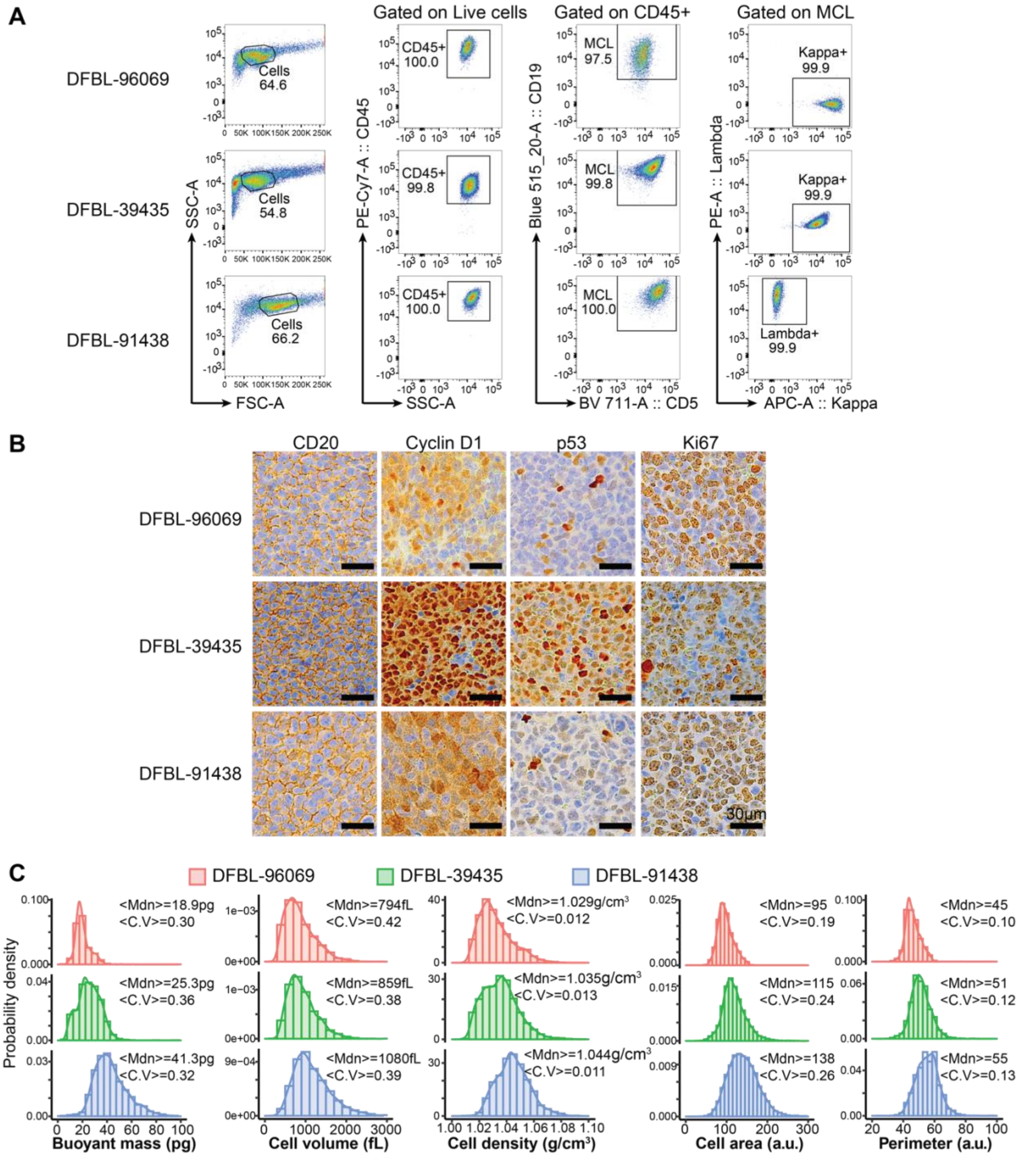


**Fig. S2. Comprehensive immunophenotypic, histopathological, and biophysical characterization of three MCL PDX models. A)** Flow cytometry analysis showing the immunophenotype of the three PDX models of MCL DFBL-96069, DFBL-39435, DFBL-91438. **B)** IHC images showing human CD20, cyclin D1, P53, and Ki67 expression in spleen cells from DFBL-96069, DFBL-39435, and DFBL-91438. All models express CD20, with strong cyclin D1 and Ki67 nuclear expression. DFBL-39435 shows strong p53 expression, while DFBL-96069 and DFBL-91438 have low TP53 levels. Images at 20X magnification, scale bar: 30 µm. **C)**

Measurement of single-cell buoyant mass, volume, and density of MCL cells from three PDX models was assessed using the fluorescence exclusion-coupled SMR. Total cell area and perimeter were evaluated with the Amnis® Imaging Flow Cytometer. The median (Mdn) and coefficient of variation (C.V) are indicated next to each graph.

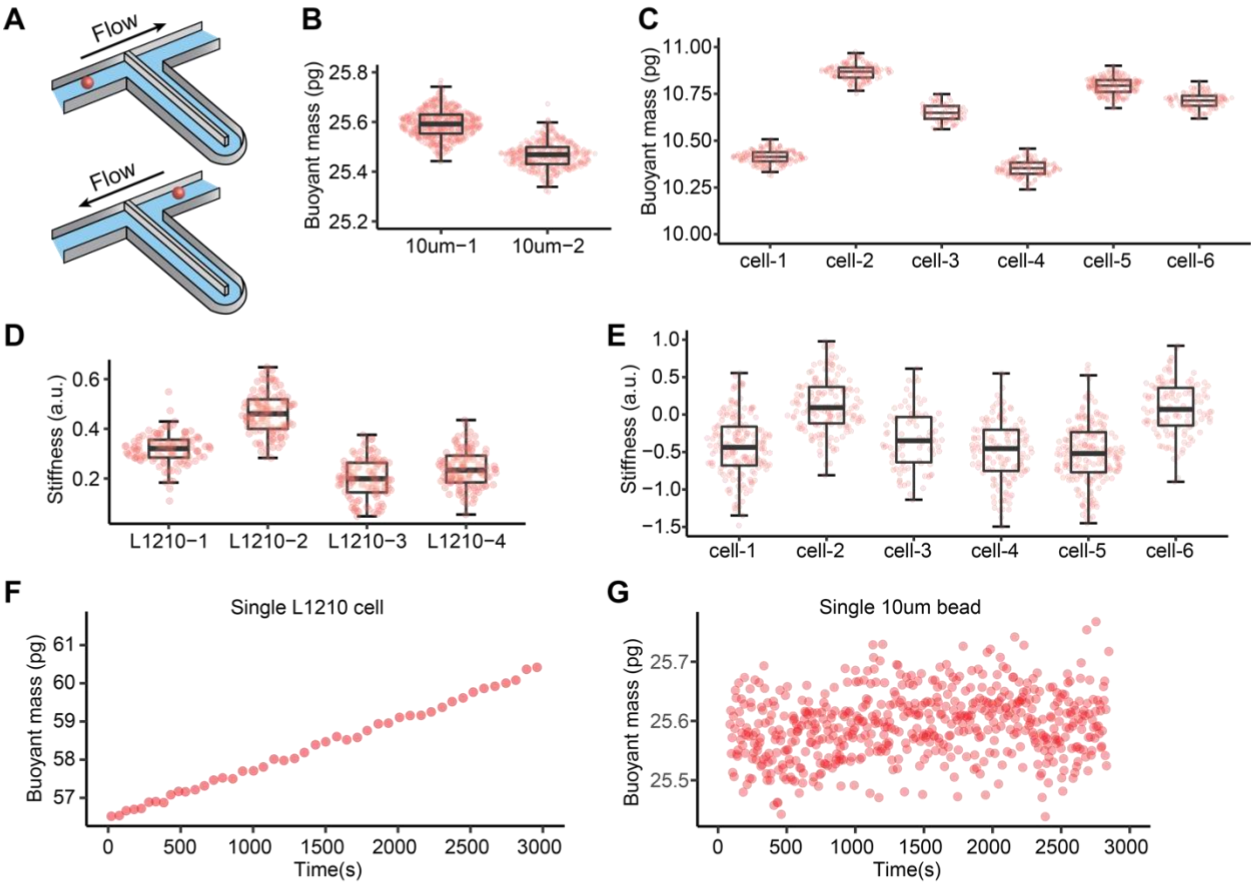


**Fig. S3. Single-cell trapping on the SMR for repeated mass and stiffness measurement of an individual cell**. **A)** Schematic of single bead/cell trapping on the SMR. Single-cell mass and stiffness is repeatedly measured as the bead/cell flows back and forth through the vibrating cantilever. To evaluate the technical interquartile ranges (IQRs) for mass measurements, we utilized 10 µm polystyrene beads **(B)** and naïve B cells isolated from human PBMCs **(C)**. To evaluate the IQRs for stiffness measurements, we used L1210 **(D)** and naïve B cells isolated from human PBMCs **(E)**. For assessing the technical IQR of mass measurements, the polystyrene bead is preferred over cell lines. This is because cell mass can vary during trapping **(F)** (*1*) —cells may accumulate mass as they grow or lose mass if apoptosis is induced —whereas the bead's mass remains constant throughout the experiment **(G)**. In contrast, when evaluating the technical IQR for stiffness measurements, cell-based samples provide a more representative range. Polystyrene beads are significantly stiffer than cells, meaning their stiffness values do not fall within the same range as those of the cells. The stiffness measurements from single-bead trapping data are available in our previous publication, Kang et al. (2019) (*2*). Additionally, since the technical IQR for stiffness differs between large and small cells, the L1210 cell line was chosen to represent larger cells (approximately 10–12 µm in diameter), while naive B cells represent smaller cells (approximately 6 µm in diameter).

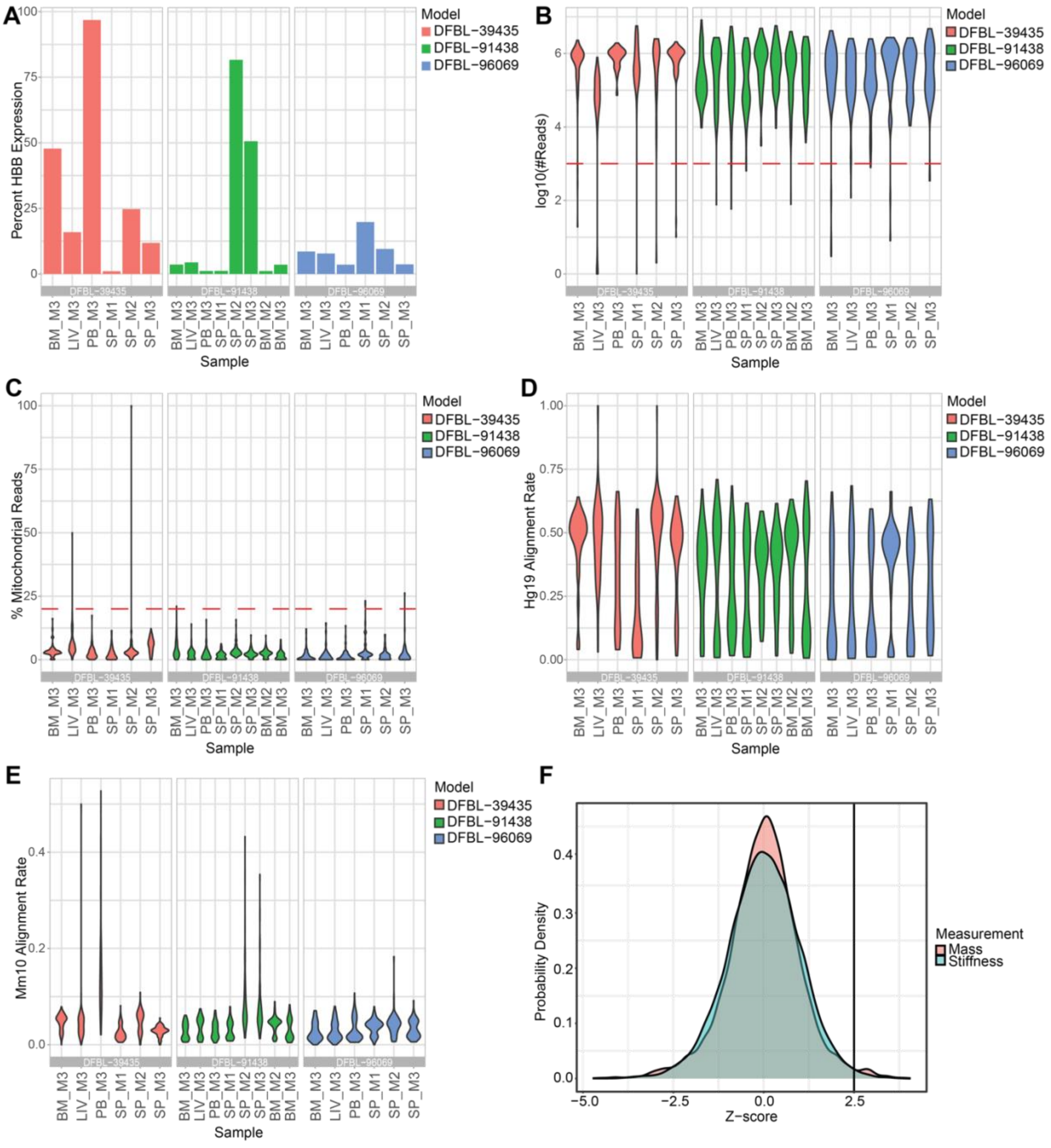


**Fig. S4. Quality control metrics of scRNA-Seq data. A)** Percentage of hemoglobin subunit beta (*HBB*) gene expression, **B)** Number of reads per cell, **C)** Percentage of mitochondrial reads, **D)** Alignment rate to human genome (Hg19), and **E)** Alignment rate to mouse genome (mm10) of human-enriched cells from DFBL-96069, DFBL-39435, DFBL-91438 isolated from different tissues (SP, spleen; PB, peripheral blood; BM, bone marrow; and LIV, liver). One to three biological replicates, each from a different mouse (M1, M2 or M3), were included per tissue. These metrics are used to assess sequencing depth, cell viability, and contamination, ensuring the integrity of the single-cell RNA sequencing data. **F)** Z-score distributions of genes correlated with cell mass and stiffness, and genes with z-score>2.5 are selected for ontology enrichment analysis.

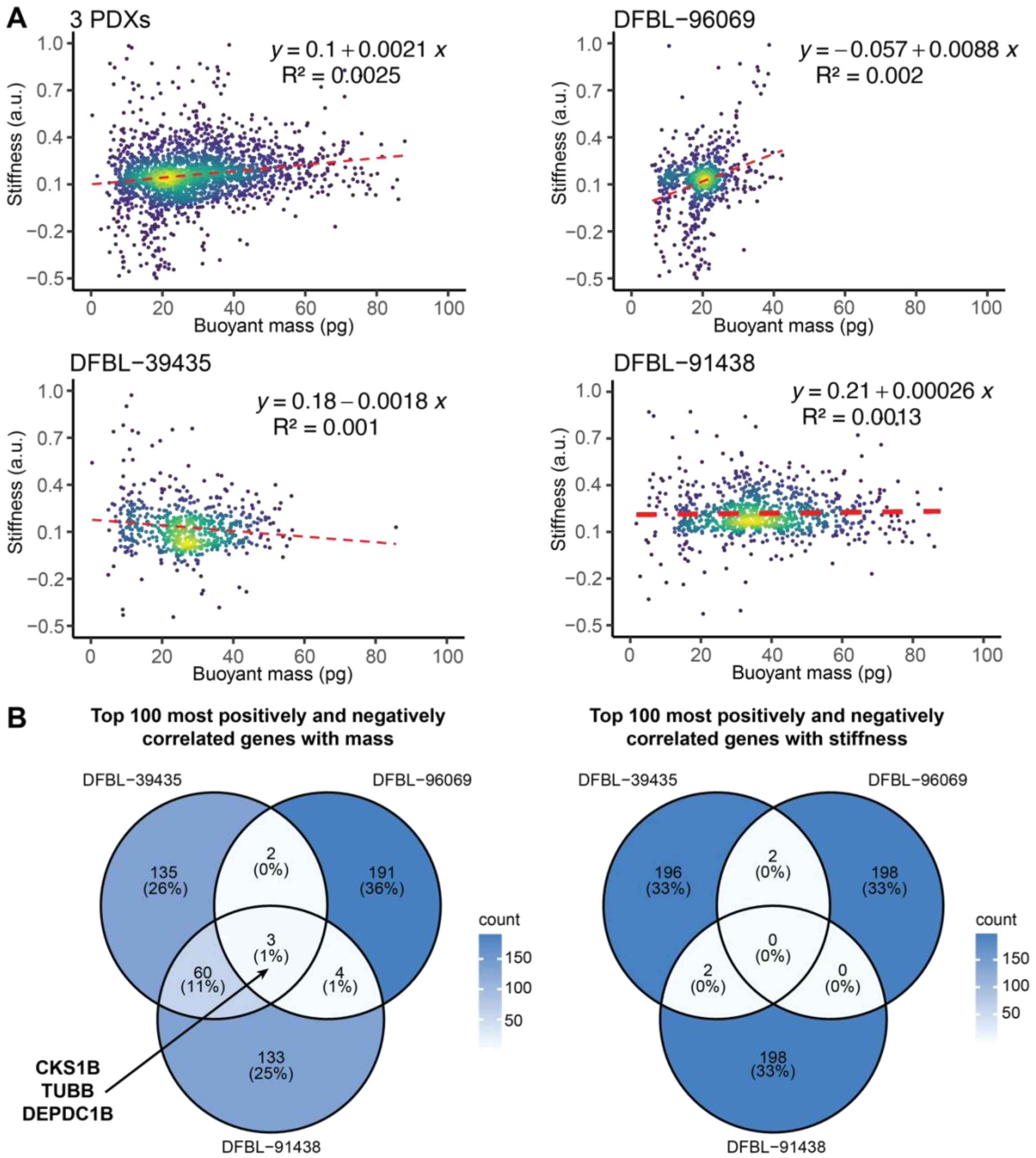


**Fig. S5. Biophysical correlation between mass and stiffness within individual MCL PDX models and their associations with gene expression. A)** Scatterplots show the relationship between buoyant mass and stiffness for individual MCL cells in three PDX models (DFBL-96069, DFBL-39435, and DFBL-91438), incorporating all cells measured from various tissues (spleen, peripheral blood, bone marrow, and liver) as shown in Figure 2. The linear regression fits (red dashed lines) and $R^2$ values indicate weak or negligible correlations within each model. Overlaid heatmaps highlight the density distributions of cells in the scatterplots. **B)** Venn diagrams show the number of shared genes across three PDX models via transcriptome-wide correlation analyses within each model, among the top 100 most positively and negatively correlated genes with mass (left) or stiffness (right).

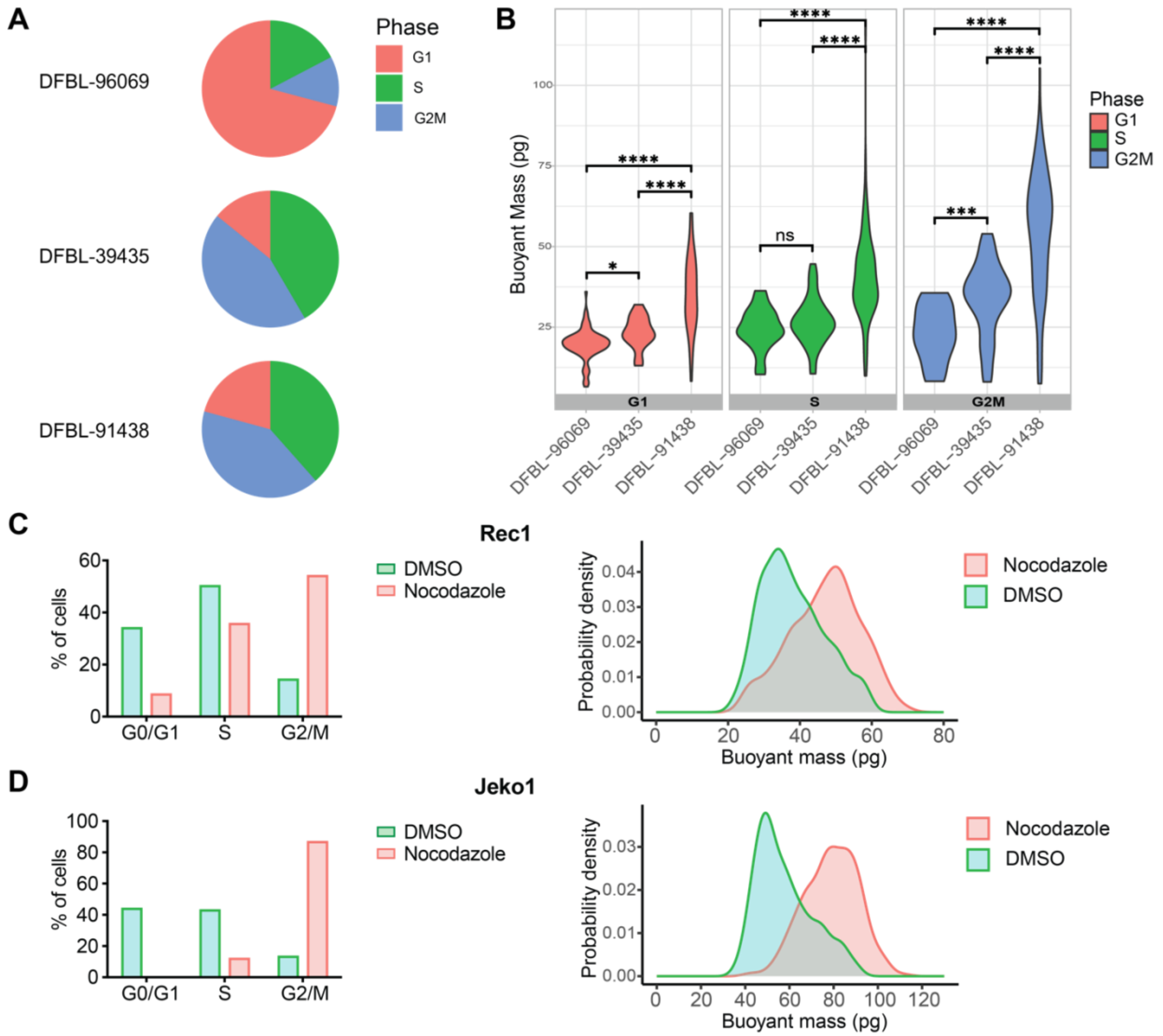


**Fig. S6. Impact of cell cycle regulation on the buoyant mass of MCL cells. A**) Pie charts summarizing the relative proportions of cells in each cell cycle phase for the three PDX models, illustrating variation in the cell cycle distribution across models. **B)** Violin plots displaying the distribution of buoyant mass for cells in different phases of the cell cycle (G0/G1, S, and G2/M) across three PDX models. C, D) Histograms showing changes in cell cycle distribution (left) by flow cytometry using PI staining and single-cell mass distribution (right) measured by the SMR after 20-hour treatment with 40 ng/mL nocodazole or DMSO in Rec1 **(C)** and Jeko1 **(D)** cells.

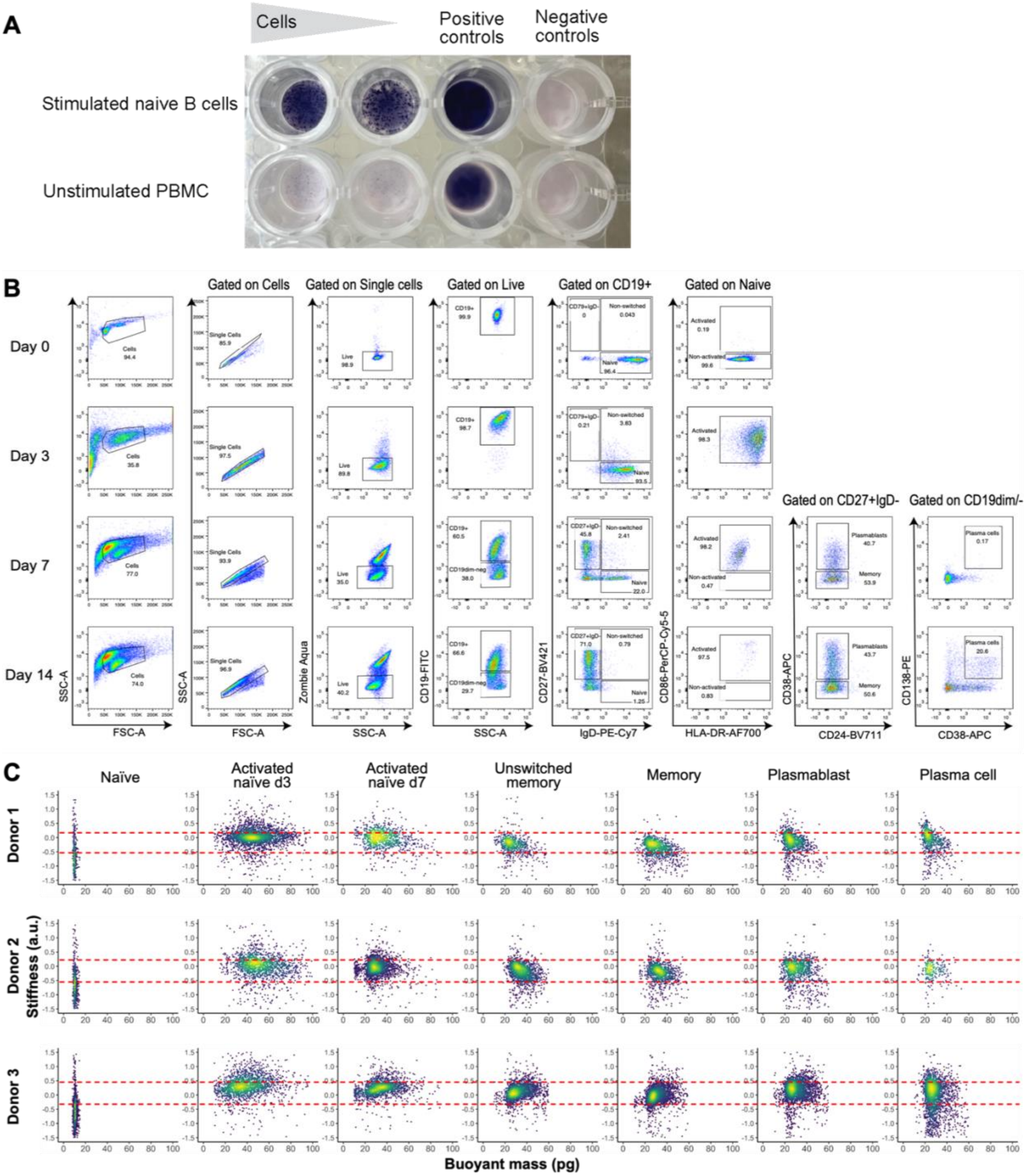


**Fig. S7. Detection of IgG secretion and biophysical profiling of B cells across differentiation stages. A)** Representative image of the ELISpot assay detecting total human IgG secretion in unstimulated PBMCs and stimulated naïve B cells cultured for seven days in RPMI + 10% FBS or the ImmunoCult™ human B cell expansion media respectively. Cells were incubated in the coated plate with RPMI + 10% FBS for 24 hours. Supernatant from the cell culture served as a positive control, while RPMI + 10% FBS medium was used as a negative control. **B)**

Representative gating strategy used to characterize human primary B-cells at various differentiation stages by flow cytometry following ex vivo activation. **C)** Buoyant mass vs stiffness profiles of single-cell B cells at different stages of differentiation isolated from three healthy donors.

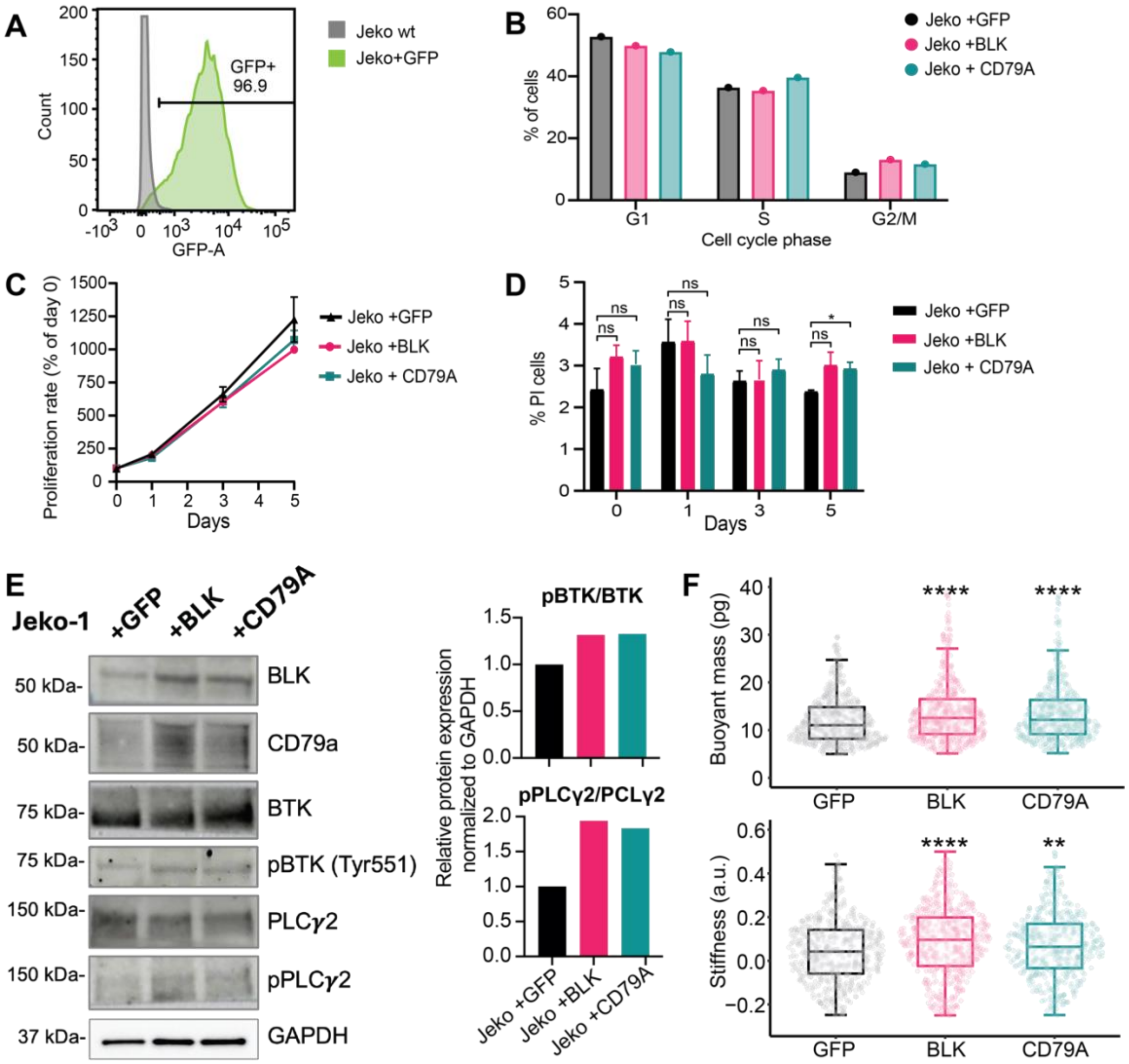


**Fig. S8. Functional analysis of BLK and CD79A overexpression in Jeko-1 cells. A)** FACS analyses of GFP staining in Jeko1$^{+GFP}$ cells in comparison to Jeko-1 wild type cells. **B)** Representative cell cycle distribution measured by flow cytometry following propidium iodine (PI) staining. **C)** Cell proliferation of Jeko-1 cells overexpressing GPF, BLK or CD79A over 5 days, as measured by flow cytometry. Data are presented as mean ± SEM of n= 3. **D)** Histograms showing the mean ± SEM of the percentage of PI positive cells measured by flow cytometry (n=3). **E)** Representative images of western blot showing downstream effectors of CD79A and BLK proteins in Jeko-1 overexpressing GFP, BLK or CD79A following LPS treatment. One representative of two western blots is shown. Quantification of the relative expression levels of phospho-BTK (pBTK/BTK) and phospho-PLCγ2 (pPLCγ2/PLCγ2) are shown on the right. **F)** Single-cell mass and stiffness measurements obtained using the SMR from >500 Jeko-1 cells overexpressing GFP, BLK or CD79A following LPS treatment. One representative experiment from two biological replicates is shown. ****P < 0.0001, **P < 0.01 as compared between indicated groups (Student's t-test).

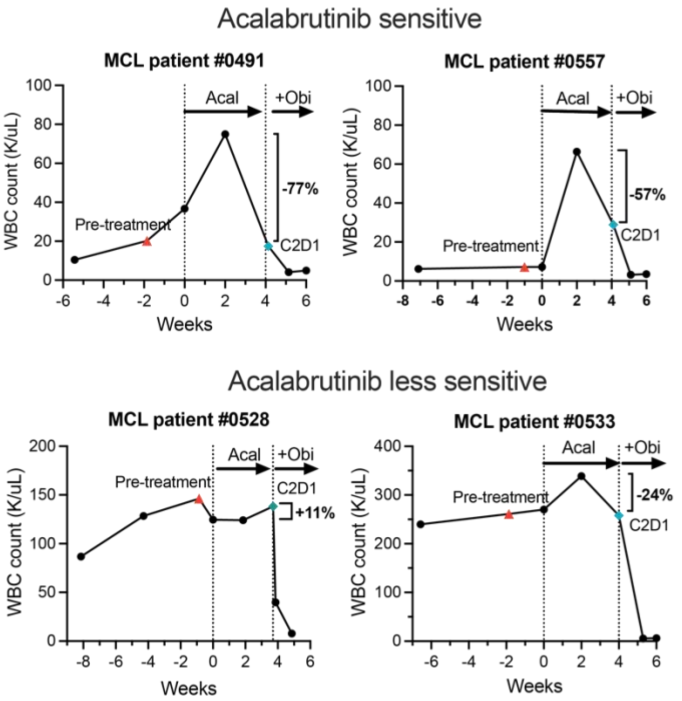


**Fig. S9. White blood cell dynamics of acalabrutinib-treated patients with MCL.** Total white blood cell (WBC) count over time in the acalabrutinib sensitive and less sensitive patient groups. The patients were treated for 4 weeks with acalabrutinib (Acal) alone before the addition of obinutuzumab (Obi). All patients except #0528 experience lymphocytosis after the start of acalabrutinib. The percent WBC reduction on Acal is indicated and was calculated from the peak level within the 30 days of Acal treatment to the end of Acal monotherapy. The pre-treatment samples (red triangle) and the Cycle 2 day 1 (C2D1; blue diamond) samples were used in our study.

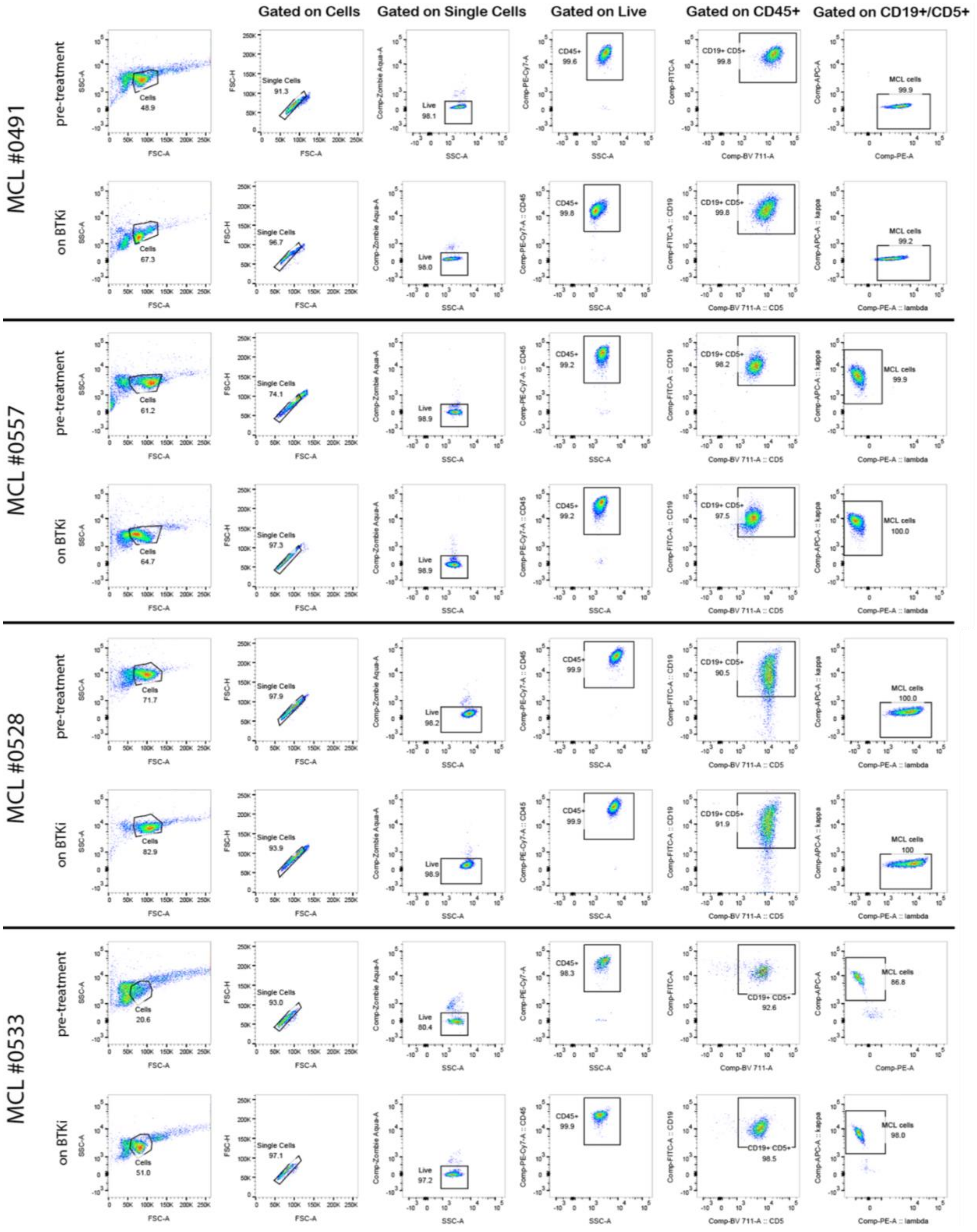


**Fig. S10. Flow cytometry of the MCL primary samples.** Flow cytometry panels used to characterize MCL tumor cells following human B cell enrichment. Cells were isolated from patients with MCL at pre-treatment or after four weeks of treatment with the BTKi acalabrutinib.

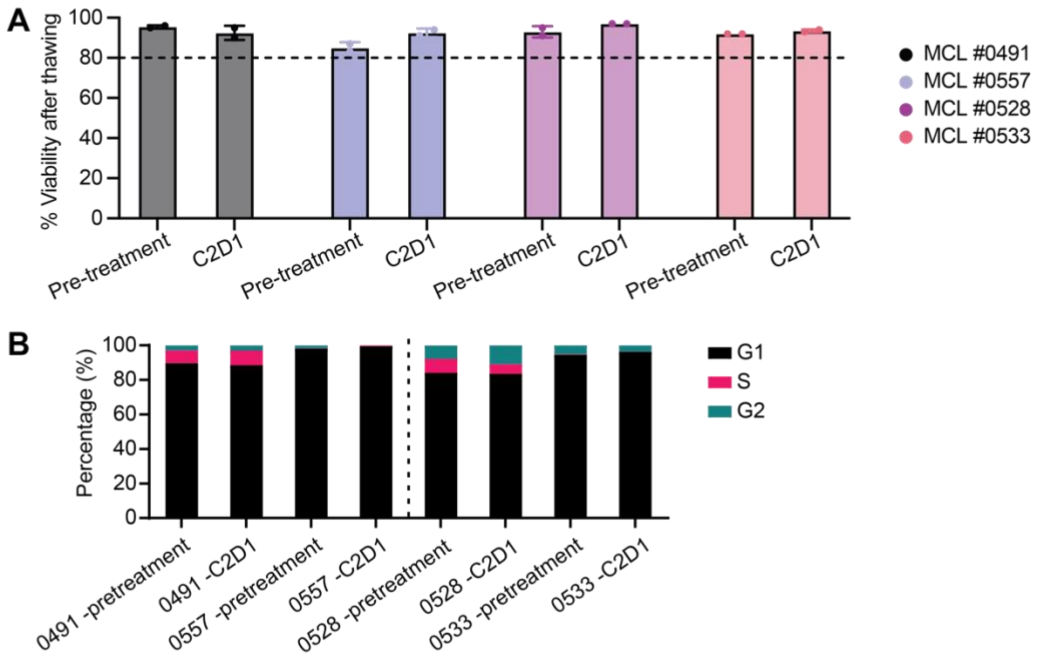


**Fig. S11. Cellular viability and cell cycle distribution in acalabrutinib-treated patients with MCL. A)** Percentage of viable cells measured by trypan blue after thawing. **B)** Cell cycle distribution of the cells analyzed by flow cytometry following propidium iodide staining after thawing.

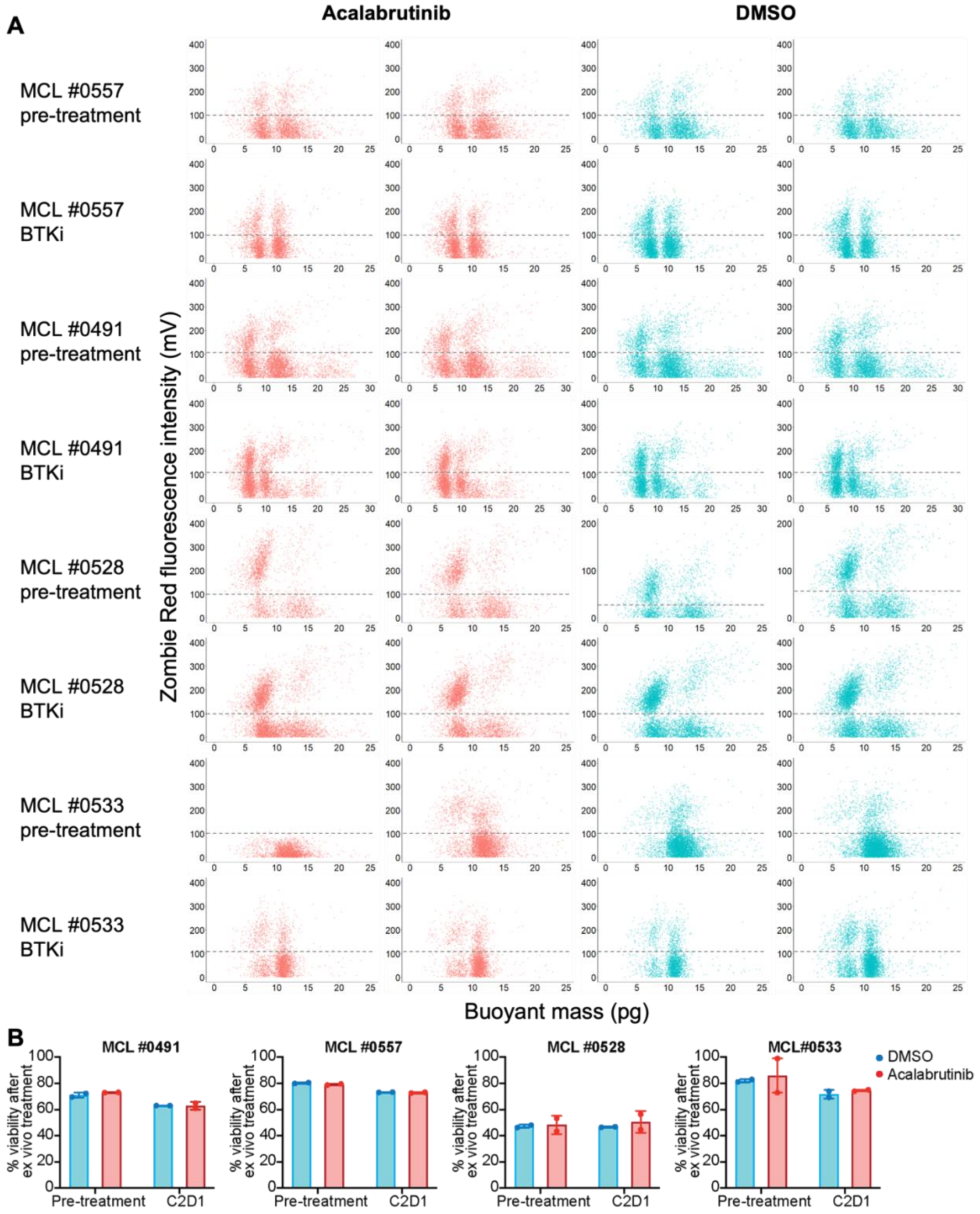


**Fig. S12. Paired mass-viability assessment of MCL cells after ex vivo drug treatment via the fluorescence exclusion-coupled SMR. A)** Paired single-cell measurement of buoyant mass (x-axis) and cell viability (indicated by ruby fluorescence intensity on the y-axis) following zombie red staining for MCL cells from AVO patient samples. Analyses were conducted on samples collected at pre-treatment and four weeks of *in vivo* treatment with the BTKi acalabrutinib. Cells were subsequently treated *ex vivo* in duplicate with either acalabrutinib or DMSO for 24 hours and analyzed using the fluorescence-coupled SMR. **B)** Percentage of viable MCL cells following *ex vivo* drug treatment, as measured by negative zombie red staining from panel A.

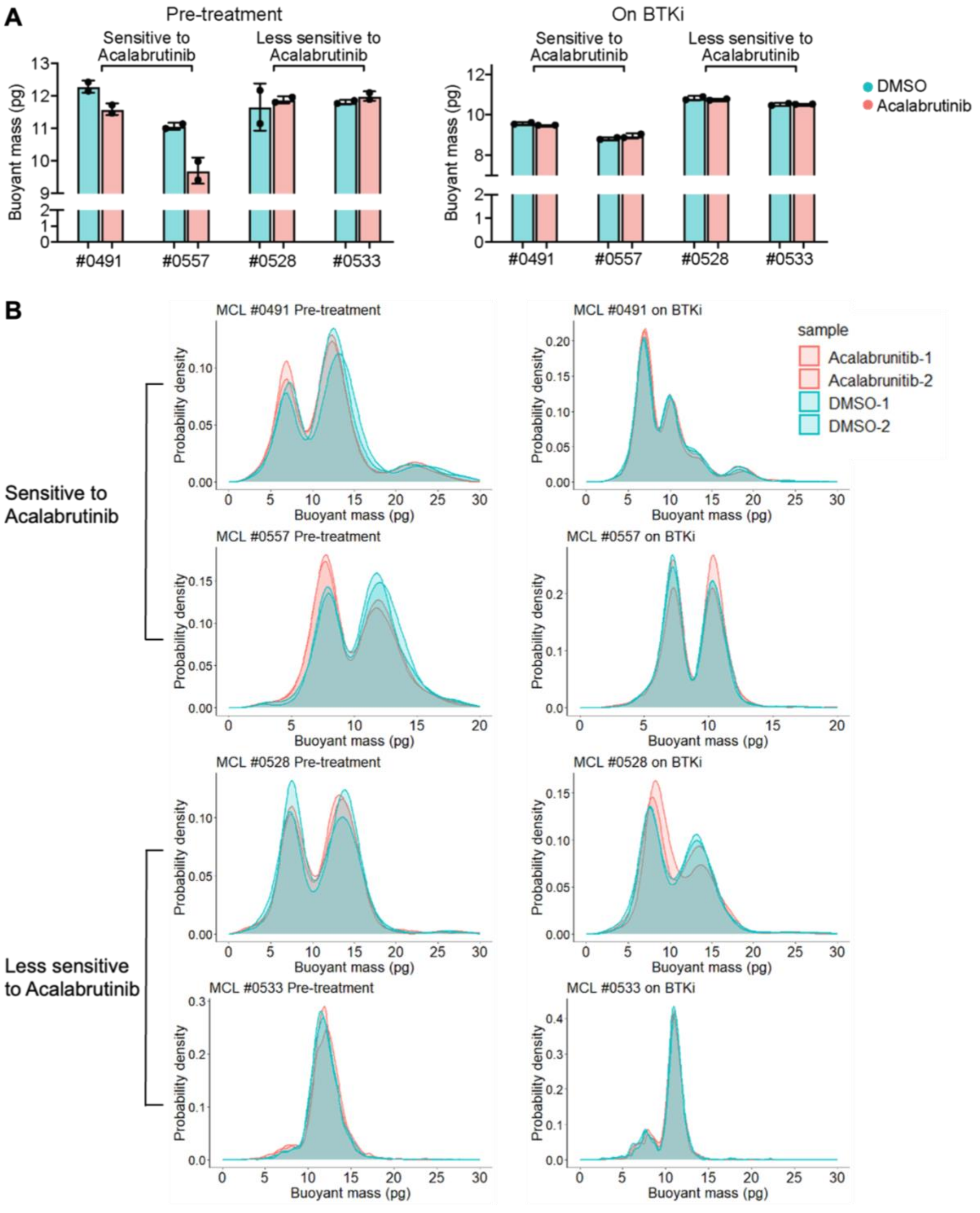


**Fig. S13. Buoyant mass profiles of MCL patient samples after ex vivo acalabrutinib treatment.** Histograms showing **(A)** the median mass or **(B)** the single-cell mass distribution of MCL primary samples of two acalabrutinib-sensitive and two less-sensitive MCL primary samples at pre-treatment (left) and after four weeks of *in vivo* treatment with acalabrutinib (on BTKi) (right). Cells were subsequently treated *ex vivo* in duplicate with either acalabrutinib or DMSO for 24 hours and analyzed using the fluorescence-coupled SMR.

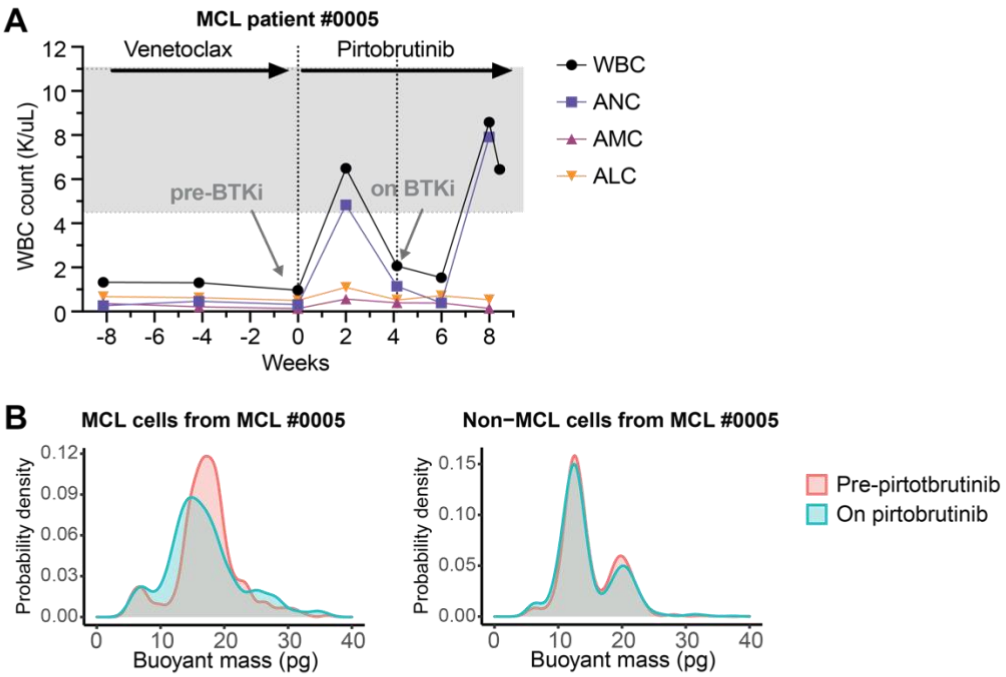


**Fig. S14. Peripheral blood dynamics and single-cell mass shifts in response to pirtobrutinib in MCL. A)** Total WBC, absolute neutrophil count (ANC), absolute monocyte count (AMC) and absolute lymphocyte count (ALC) over time in a MCL patient pre- and post- pirtobrutinib treatment. At the pre-pirtobrutinib timepoint, strong evidence of active disease progression led to a change in therapy. After one cycle of pirtobrutinib, the patient exhibited signs of a transient clinical response, including recovery of neutrophils, as well as symptomatic improvements with weight gain, increased energy, and enhanced appetite. **B)** Single-mass measurements of serial peripheral blood specimens from a patient with relapsed/refractory MCL collected at pre- and post-BTKi following four weeks of pirtobrutinib treatment. MCL and non-MCL cells were enriched using FACS.

**A**

| SAMPLE ID | SAMPLE TYPE | AGE AT SAMPLE | SEX | CYTOGENETICS | IGHV | *TP53* | PREVIOUS TREATMENTS BEFORE ACALABRUTINIB |
|---|---|---|---|---|---|---|---|
| **JB-0215 PRE-TREATMENT** | PBMC | 66 | M | 90.5% 13q | unmutated | Wild type | BR+ Lenalinomide |
| **JB-0215 PROGRESSION** | PBMC | 70 | M | | unmutated | Mutated | |
| **JB-0375 PRE-TREATMENT** | PBMC | 50 | M | 34.5% 11q deletion, 20.5% 13q deletion, 19% 17p deletion | unmutated | Wild type | FCR |
| **JB-0375 PROGRESSION** | PBMC | 54 | M | | | Wild type | |
| **JB-0097 PRE-TREATMENT** | PBMC | 52 | M | 94% 11q del, 86% 13q del | unmutated | Wild type | FR, Rituxan, Spebrutinib, Ibrutinib |
| **JB-0097 PROGRESSION** | PBMC | 57 | M | | unmutated | Wild type | |

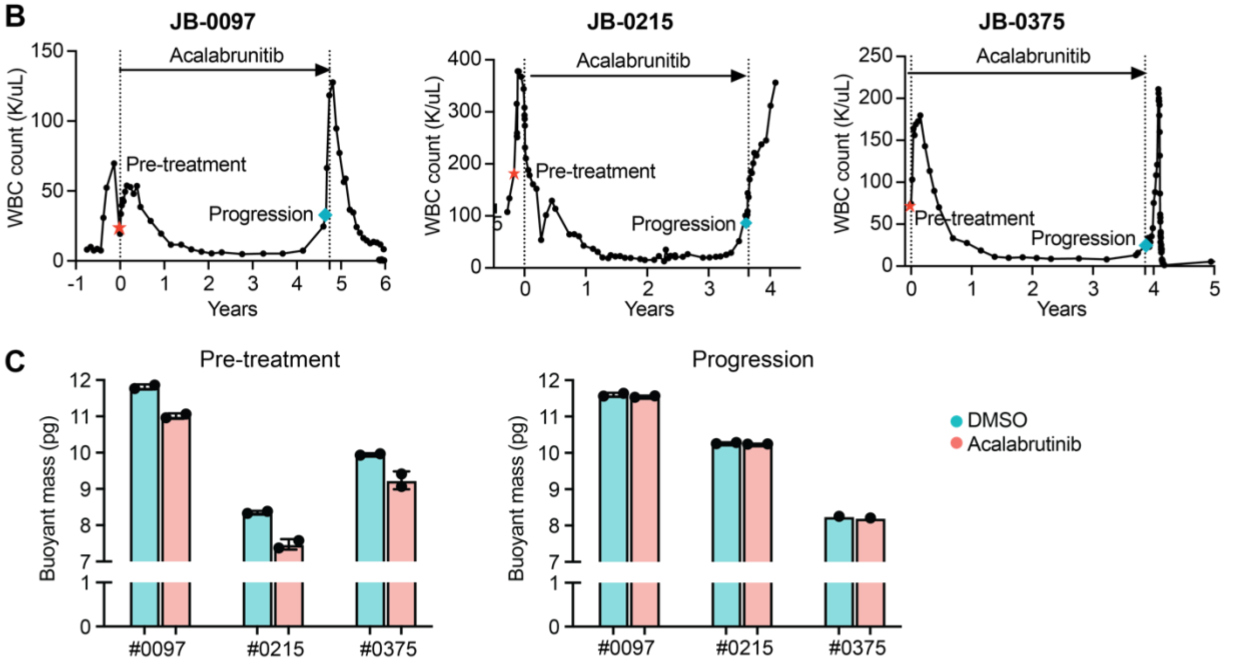


**Fig. S15. Clinical information and biophysical analysis of CLL patient samples treated with acalabrutinib. A)** Clinical and molecular characteristics of CLL patient samples. Bendamustine (B), rituximab (R), fludarabine (F), cyclophosphamide (C). **B)** WBC counts over time in CLL patients treated with acalabrutinib monotherapy. The pre-treatment (red triangle) and progression (blue diamond) samples were used in our study. **C)** Bar graphs showing the median mass measured by the SMR of three CLL primary samples at pre-treatment and progression timepoints, followed by 24 hours of ex vivo drug treatment in duplicate with DMSO or acalabrutinib.

**Table S1. Mutational landscape of three MCL PDX models identified by hybrid capture, target enrichment next generation sequencing of coding exons of 205 genes. (separate file)**

https://www.dropbox.com/scl/fi/21lfv71c9x6kyyltleflw/Table-supp1_HemoSeq-mutations-PDX_clean.xlsx?rlkey=y20xxtyxw73ocebp3cs8ky2cx&st=4mwepsnq&dl=0

**Table S2. Gene ontology results from GSEA analysis of top genes (z-score > 2.5) correlated with mass and stiffness across DFBL-39435, DFBL-96069, and DFBL-91438. (separate file)**

https://www.dropbox.com/scl/fi/koydx4ijyarvqsohxwn95/TableS2-full-gene-ontology-results.xlsx?rlkey=nv6saw21ff2t55kvwqpiey43h&st=fly1keq0&dl=0

**Table S3. List of genes correlating with cell mass, cell stiffness and both mass and stiffness across DFBL-39435, DFBL-96069 and DFBL-91438. (separate file)**

https://www.dropbox.com/scl/fi/3trn9kf7ebospm9jzrn6s/TableS3_correlation-gene-list.xlsx?rlkey=hbs77c5jtcidmpe7tysd67ri1&st=7i2583qn&dl=0

**Table S4. MCL patient's clinical information.**

| Sample ID | #0491 | #0557 | #0528 | #0533 |
|---|---|---|---|---|
| **Sex-Age** | M-69 | M-68 | M-69 | M-70 |
| **Prior treatment for MCL** | Induction BR/R-AraC, maintenance R | None | None | None |
| **Prior ASCT** | Yes | No | No | No |
| **MCL stage** | III | IV | IV | III |
| **ECOG Performance Status** | 1 | 1 | 1 | 0 |
| **Histological subtype** | Classic | Classic | Classic | Classic |
| **Bone marrow involvement** | 51% | 7% | 50% | 76% |
| **Ki-67 index** | 15-20% | <30% | 40% | Not available |
| **TP53 status** | Mutated | Wild-type | Mutated | Mutated |
| **Cytogenetics** | 46,XY,del(1)(p22p13), add(4)(q21), -10,t(11;14)(q13; q32), add(13)(q22),del(14)(q24q32)(q24q32),+mar[8] /46,idem,t(5;15)(q13;q22), add(10)(p13)[6]/46,XY[6] .nuc ish(CCND1,IGH)x3(CCND1 con IGHx2-3)[134/200] | 46,XY,t(11;14)(q13;q32)[4]/ 46,idem,del(9)(q12q3?4)[cp5]/46, XY[11] | 43,-Y,add(X)(p11.2),-9 ,-10,-11,t(11;14)(q13;q32),add(12)(p1?2),-13,-18,+2-4mar[cp4]/45,X,Y[4]/46, XY[12] | 45,XY,t(11;14)(q13;q32), der(15;17)(q10;q10)[4]/45, idem,t(2;13)(p13;q12), -12, +mar[cp11]/46, XY[cp5] |
| **Pathogenic known mutations** | **PPM1D:** c.1528_1529insA (p.N512Kfs*16)7.1% VAF, c.1445delT(p.L482Rfs*3) 1% VAF; **TP53**: c.638G>A (p.R213Q) 66.2% VAF; | **NOTCH1**: c.7541_7542delCT (p.P2514Rfs*4) 5.3% VAF | **TP53:** c.524G>A (p.Rl75H) 95% VAF **ATM:** c.901+1G>C (splice site) 26.4% VAF | **CCND1**: c.130T>G (p.Y44D) 38.5% VAF; **TP53**: c.743G>A (p.R248Q) 65.7% VAF |

| | | | | |
|---|---|---|---|---|
| | **ZRSR2**: c.1070_ 1106delTTGGGAAGAA CTCCGAAAGGAGGGAGAGGATGGGCCAinsGC(p.F357Cfs*16) 7.3% VAF | | | |
| **Copy number variations** | gain **SF3B1**, **IDH1** (on 2q); CN-LOH **TP53** (on 17p) | None detected | gain **RAD21**, **MYC** (on 8q); 1 copy deletion **JAK2**, **CDKN2A**, **CDKN2B** (on 9p); 1 copy deletion **ABL1**, **NOTCH1** (on 9q); gain **WT1** (on 11p); gain **ATM**, amplification **KMT2A**, 1 copy deletion **CBL** (on 11q); 1 copy deletion **ETV6**, **ETNK1**, **KRAS** (on 12p); 1 copy deletion **FLT3** (on 13q); gain **MAP2K1**, **IDH2** (on 15q); CN LOH **TP53** (on 17p) | 1 copy deletion **DNMT3A** (on 2p), **PRPF8** (on 17p), **TP53** (on 17p) |
| **Variants of unknown significance** | **MPL**: c.655C>G (p.Q219E) 58.7% VAF | **CDKN2A**: c.106G>A (p.A36T) 41.7% VAF | **ATM:** c.8293G>A (p.G2765S) 41.2% VAF; **CREBBP:** c.1369A>G (p.I457V) 48% VAF | **TET2**: c.4787A>G (p.N1596S) 48.3% VAF |

**Table S5. Antibodies used for flow cytometry.**

| Antibody | Clone | Catalog # | Assay(s) |
|---|---|---|---|
| Pe/Cy7 anti-human CD45RA | HI100 | BioLegend, #304125 | PDX and primary cell characterization |
| BV711 anti-human CD5 | L17F12 | BD Biosciences, #742552 | PDX and primary cell characterization |
| APC anti-human Kappa | TB28-2 | BD Biosciences, #341098 | PDX and primary cell characterization |
| PE anti-human Lambda | 1-155-2 | BD Biosciences, #642919 | PDX and primary cell characterization |
| BB515 anti-human CD19 | HIB19 | BD Biosciences, #564456 | PDX and primary cell characterization |
| APC anti-mouse CD45 | 30-F11 | BioLegend, #103112 | PDX and primary cell characterization |
| BV421 anti-human CD27 | M-T271 | BD Biosciences, #562513 | B cell characterization |
| APC anti-human CD38 | HIT2 | BD Biosciences, #560980 | B cell characterization |
| Pe-Cy7 anti-human IgD | IA6-2 | BD Biosciences, #561314 | B cell characterization |
| BV711 anti-human CD24 | ML5 | BD Biosciences, #563401 | B cell characterization |
| PE anti-human CD138 | MI15 | BD Biosciences, #561704 | B cell characterization |
| PerCP-Cy5.5 anti-human CD86 | 2331 | BD Biosciences, #561129 | B cell characterization |
| PE anti-human CD86 | IT2.2 | BD Biosciences, #555665 | B cell characterization |
| AF700 anti-human HLA-DR | G46-6 | BD Biosciences, # 560743 | B cell characterization |
| Zombie Aqua Fixable Viability Kit | | BioLegend, #423102 | PDX, primary cell and B cells characterization |
